# Regulation of neuroinflammation by astrocyte-derived cholesterol

**DOI:** 10.1101/2022.12.12.520161

**Authors:** Hao Wang, Joshua A. Kulas, Yang Yu, Joshua L. Milstein, Claire E. Peng, Holden Higginbotham, Jed Robert Christensen, Michael A. Kovacs, Anton M. Schüle, Denys Oliinyk, Thierry M. Nordmann, Matthias Mann, Adriano Aguzzi, Heather A. Ferris, Scott B. Hansen

**Affiliations:** Department of Molecular Medicine, The Scripps Research Institute, Jupiter, FL 33458, USA; Department of Neuroscience, The Scripps Research Institute, Jupiter, FL 33458, USA; Skaggs Graduate School of Chemical and Biological Sciences, The Scripps Research Institute, Jupiter, FL 33458, USA; Institute for the Science of the Aging Brain (ISAB), St. Gallen, Switzerland; Division of Endocrinology and Metabolism, University of Virginia, Charlottesville, VA 22908, USA; Center for Brain Immunology and Glia, Department of Neuroscience, University of Virginia, Charlottesville, VA 22908, USA; Chinese Institutes for Medical Research (CIMR), Institute for Medical Physiology, Beijing China; Capital Medical University, Beijing China; Department of Biology, Brigham Young University-Idaho, Rexburg, ID 83460, USA; Department of Proteomics and Signal Transduction, Max-Planck Institute of Biochemistry, Martinsried, Germany; Center for Membrane and Cell Physiology, University of Virginia, Charlottesville, VA 22908, USA

## Abstract

Neurodegeneration and its concomitant loss of cognitive function are closely linked to neuroinflammation and lipid accumulation, particularly cholesterol. In the brain, astrocytes synthesize cholesterol and deliver it to surrounding cells via apolipoprotein E (apoE). Astrocyte activation increases apoE secretion and promotes neuronal cell death, yet the link between astrocyte-derived cholesterol and inflammation remains unclear. Here we show that pro-inflammatory cytokines stimulate cholesterol synthesis and release from astrocytes. Immune cells take up this cholesterol, which clusters pro-inflammatory receptors in lipid rafts, amplifying inflammatory signaling and gene expression. Astrocyte-specific knockout of cholesterol synthesis (via SREBP2 deletion) reduces inflammatory cytokine production in an Alzheimer’s disease (AD) mouse model and shortens neuroinflammation triggered by peripheral lipopolysaccharide (LPS) injection. These findings identify astrocyte-derived cholesterol as a paracrine signal that helps drive neuroinflammation in microglia and brain-resident macrophages.

## Introduction

Neurodegeneration is characterized by a progressive loss of cognitive function and is strongly associated with aging and chronic neuroinflammation ^1,2^. In healthy brains, inflammation is typically acute and self-resolving, but it can become chronic with age, disease, or injury^3,4^. Chronic neuroinflammation is a hallmark of prominent neurodegenerative disorders, including Alzheimer’s disease (AD) and Parkinson’s disease^1,2,5,6^.

At the cellular level, neuroinflammation arises from interactions between microglia and astrocytes^7,8^. Microglia, the brain’s primary immune cells release cytokines that activate astrocytes^7,9^ (**Figure S1A**). Astrocytes, in turn, provide metabolic and structural support to neurons and release both pro- and anti-inflammatory factors that modulate microglia activity^7,10^. Astrocytes also produce most of the cholesterol in the brain, independent of peripheral sources ^11,12^.

Upon microglial activation, astrocytes become reactive and increase secretion of saturated long-chain fatty acids and apolipoprotein (apoE)^9,13^. Conditioned medium from activated astrocytes induces cytokine release from microglia^14^, potentially creating a self-reinforcing inflammatory cycle^12^. However, the specific signals linking astrocyte cholesterol production to microglial inflammation are not fully defined.

Because apoE primarily transports cholesterol and is upregulated in reactive astrocytes, we hypothesized that astrocyte-derived cholesterol acts as a bioactive signal that escalates neuroinflammation. Supporting this idea, astrocyte-derived cholesterol is paracrine signal escalating amyloid production in AD mouse neurons^15^. Selectively knocking out a key cholesterol synthesis protein, sterol regulatory element-binding protein 2 (SREBP2), from astrocytes abolished amyloid production and tau phosphorylation in a familial AD (FAD) mouse model^15^.

Here we demonstrate that pro-inflammatory cytokines drive cholesterol synthesis and secretion from astrocytes, and that astrocyte-derived cholesterol promotes inflammatory gene expression in immune cells. *In vivo*, loss of astrocyte cholesterol reduces neuroinflammation in AD mice and accelerates resolution of inflammation after peripheral LPS challenge. Thus, astrocyte cholesterol links amyloid pathology to neuroinflammation and represents a potential therapeutic target.

## Results

### Pro-inflammatory cytokines stimulate cholesterol production in astrocytes

Interleukin 6 (IL6) and tumor necrosis factor alpha (TNFα) are key cytokines that activate astrocytes and stimulate cholesterol synthesis in hepatocytes ^7–9^, the primary cholesterol-producing cells of the periphery (**Figure S1B**)^16^. We tested whether they similarly increase cholesterol production in primary mouse astrocytes. Astrocytes were treated with cytokines for 24h, the cytokines removed, and the cholesterol accumulation measured in the astrocyte-conditioned media (ACM).

We found IL6 and TNFα increased both total and free cholesterol levels in ACM by 25 to 40% (**Figure 1A-B**). In contrast, free cholesterol levels in the cell lysates remained unchanged following either treatment (**Figure 1C**), suggesting the cultured cells maintain a relatively constant intracellular concentration of free cholesterol while increasing cholesterol release into the extracellular space. Total cholesterol in the cell lysates (free and esterified) remained constant for IL6 treatment (**Figure 1D**) but increased slightly for TNFα and lipopolysaccharide (LPS) treatment. In contrast, the same cytokines did not increase cholesterol release from primary microglia or N2a neurons (**Figure 1E-F**), consistent with astrocytes being the primary source of brain cholesterol.

**Figure 1.**
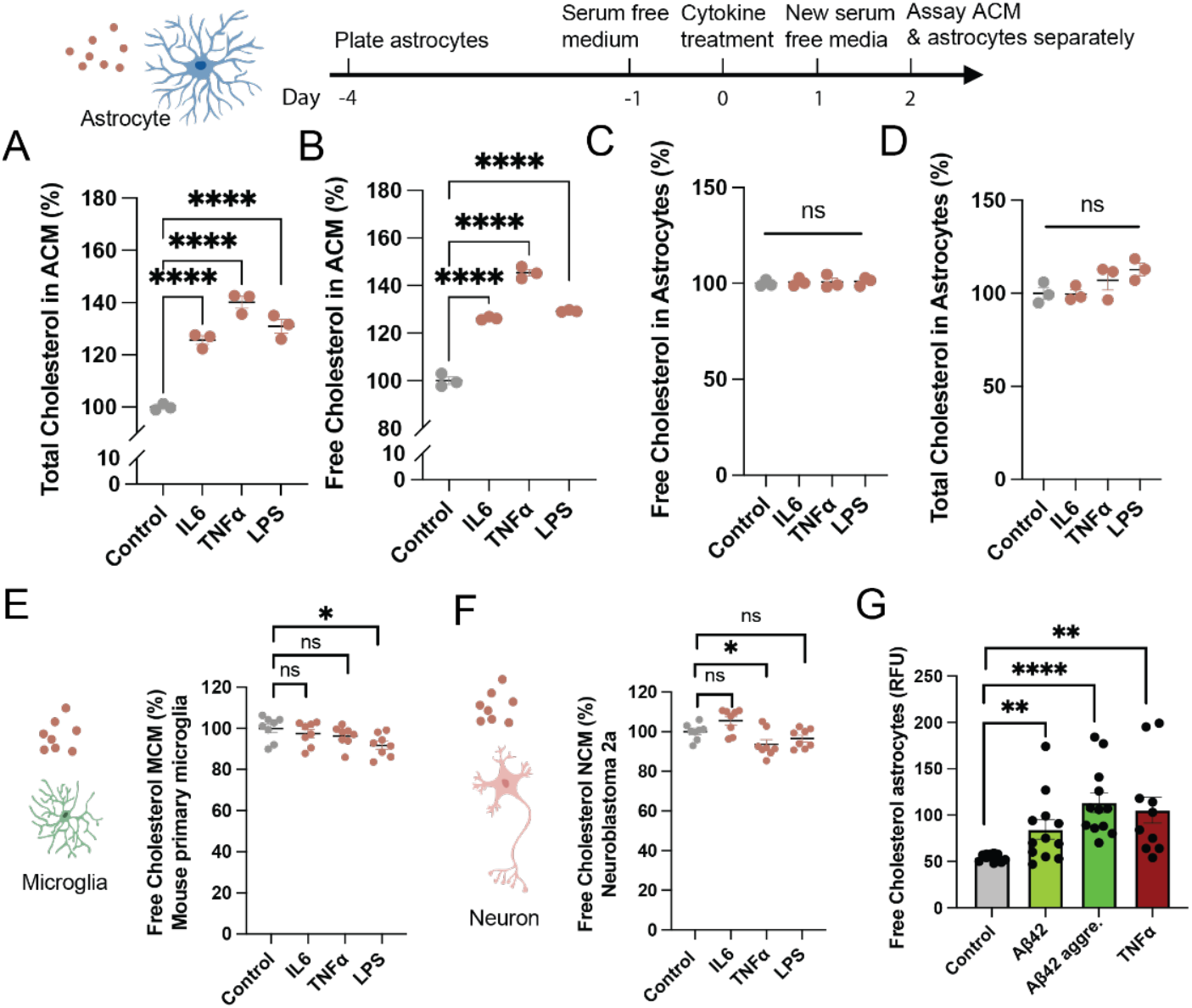
Inflammatory cytokines induce cholesterol release from astrocytes but not immune cells or neurons. (A-B) Cholesterol released into the astrocyte conditioned media (ACM) was increased by cytokines and lipopolysaccharide (LPS). Free cholesterol released into the media was measured by a fluorescent cholesterol oxidase assay. Total cholesterol is both free cholesterol and cholesterol esters. (C-D) Cholesterol in the astrocytes was measured separate from the media. (E) Cholesterol released into microglial condition media (MCM) after 100 ng/mL cytokine treatment. (F) Cholesterol released into neuronal conditioned media (NCM) after 100 ng/mL cytokine treatment. (G) Comparison of cholesterol release from primary astrocytes after application of amyloid beta 42 (Aβ42), aggregated Aβ42, and 100 ng/mL TNFα. Values are expressed as mean (each experimental point is from a well of cells (n=3, derived from 2 animals). Comparison made with a Student’s T test; *p≤0.05, ** p≤0.01, **** p≤0.0001

Dose-response experiments confirmed rapid (within 2h), cytokine-dependent increases in extracellular free cholesterol from live primary mouse astrocytes (**Figure S2A-C**). TNFα (10-100 ng/mL), IL6 (1-50 ng/mL), and Interlukin-1β (IL-1β, 1-50 ng/mL) were added to astrocyte media and the free cholesterol determined using a cholesterol oxidase assay adapted from a live choline oxidase assay (see methods)^17^. IL6 had the greatest effect, increasing total produced cholesterol by >200% over the 2 h incubation, followed by TNFα (~180%). IL-1β induced a small (20%), but also statistically significant dose dependent increase in free cholesterol. A similar treatment in HMC3 microglia showed no such change in cholesterol (**Figure S2D**). Amyloid-β (both soluble and aggregated forms) also stimulated cholesterol release from primary astrocytes, with the aggregated Aβ42 comparable to TNFα (100 ng/mL) (**Figure 1G**).

To determine whether inflammatory signaling broadly activates cholesterol biosynthetic pathways in astrocytes, we reanalyzed a recently published proteomics dataset of human iPSC-derived astrocytes stimulated with the pro-inflammatory cytokine cocktail TNFα, IL-1α, and C1q^18^. Consistent with our cholesterol measurements in primary astrocytes, multiple enzymes in the cholesterol synthesis pathway were increased following cytokine stimulation, including HMGCS1 and HMGCR, which catalyze the initial steps of cholesterol synthesis (**Figure S3**). Additional enzymes throughout the mevalonate and sterol synthesis pathways were similarly elevated. These findings from an independent dataset support cytokine-induced activation of cholesterol biosynthesis in astrocytes.

### Cholesterol induced changes in immune cell gene expression

To test whether astrocyte-derived cholesterol promotes inflammation, we analyzed RNA-seq datasets from cholesterol-treated (2 h) RAW264.7 macrophages (n=3) and HMC3 microglia-like cells (n=5) (**Figure S4A-B**). Reactome pathway enrichment analysis of differentially expressed genes (DEGs) revealed that approximately half (10/20 for RAW264.7 and 8/20 for HMC3) of the top 20 enriched pathways in both cell types were related to inflammatory or immune signaling, indicating that cholesterol broadly activates inflammatory transcriptional responses in both macrophage- and microglia-like cells (**Figure 2A-B**). The treatment itself had no effect on cell viability in a CCK8 assay (**Figure S5A-B**).

**Figure 2.**
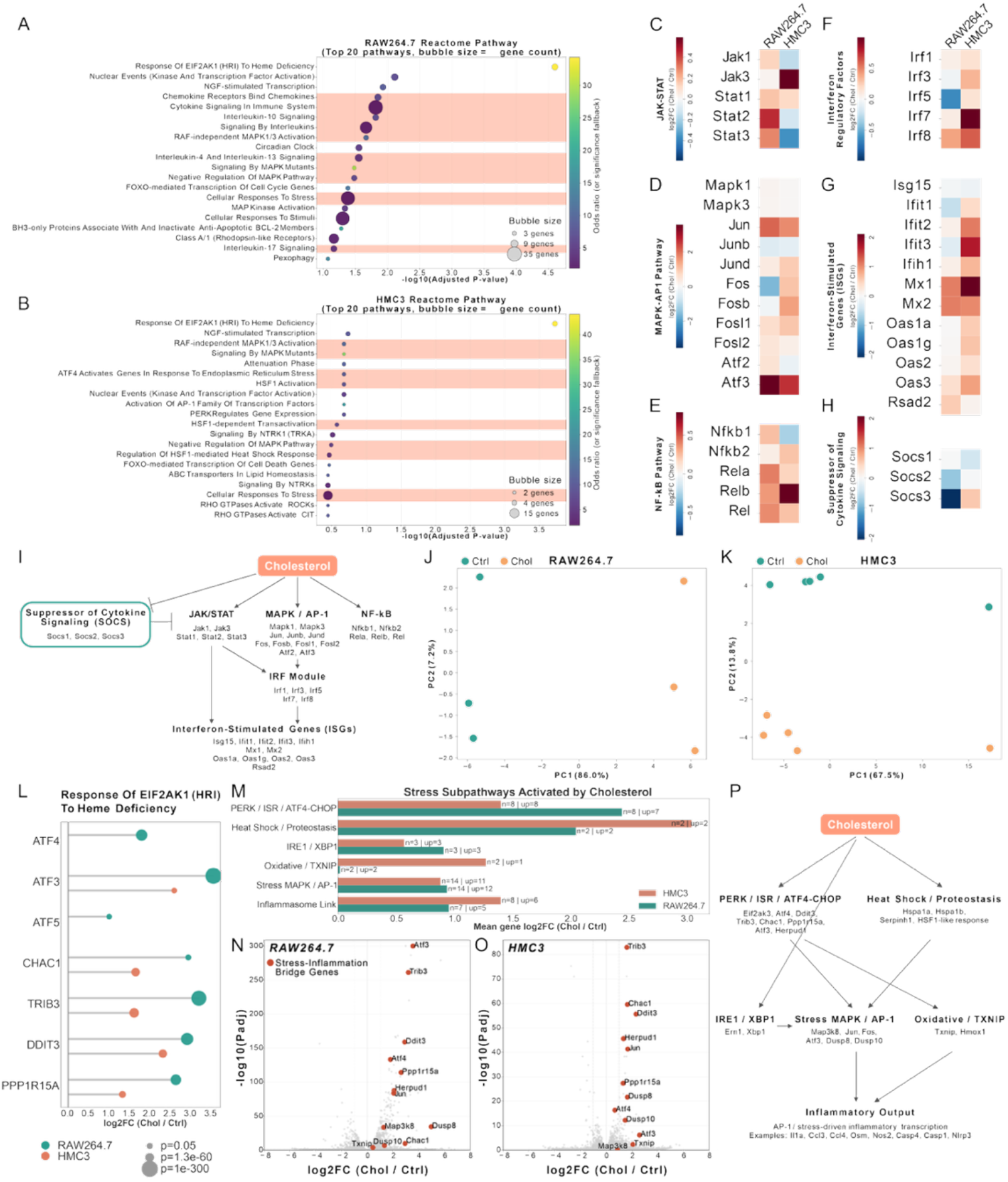
Cholesterol induces inflammatory and stress-associated transcriptional programs in RAW264.7 and HMC3 cells. (A-B) Reactome pathway enrichment analysis of differentially expressed genes (DEGs) from cholesterol-treated (chol) versus control-treated (Ctrl) RAW264.7 and HMC3 cells. Among the top 20 enriched Reactome pathways in each cell line, approximately half were related to inflammatory responses. (C-H) Heatmaps showing log2FC (chol/control) of key genes in major inflammatory signaling pathways, including JAK/STAT, MAPK/AP-1, NF-kB, interferon, and SOCS. The first column shows results from RAW264.7 cells and the second column shows results from HMC3 cells. (I) Schematic model summarizing the inflammatory pathways induced by cholesterol, including JAK/STAT, MAPK/AP-1, and NF-kB, together with reduced expression of SOCS genes, which normally inhibit JAK/STAT signaling. (J-K) Principal component analysis (PCA) performed using the transcriptional profiles of the selected inflammatory pathway genes. Cholesterol-treated and control samples separated clearly in both RAW264.7 and HMC3, indicating a distinct inflammation-associated transcriptional response. (L) Bubble plot showing the seven genes contributing to enrichment of the Reactome pathway Response Of EIF2AK1 (HRI) To Heme Deficiency, the strongest enriched Reactome pathway in both RAW264.7 and HMC3 DEG analyses. The x-axis shows log2FC (chol/control), and bubble size represents adjusted p-value. Green indicates RAW264.7 and salmon indicates HMC3. (M) Summary plot of six stress-signaling subpathways. The x-axis shows the mean gene log2FC (chol/control) for each stress subpathway. For each bar, the total number of genes in the pathway (n) and the number of genes upregulated by cholesterol (up) are indicated. Green indicates RAW264.7 and salmon indicates HMC3. (N-O) Volcano plots for RAW264.7 and HMC3 highlighting genes proposed to bridge stress signaling and inflammation. All highlighted bridge genes were upregulated in response to cholesterol, and several were among the most strongly induced DEGs. (P) Schematic model of a cholesterol-triggered stress-to-inflammation mechanism, in which cholesterol activates stress pathways that in turn promote inflammatory transcriptional responses.

To further define the pathways altered by cholesterol, we examined the transcriptional behavior of key genes in major inflammatory signaling pathways, including JAK/STAT, MAPK/AP-1, NF-kB, interferon-related pathways, and Suppressor of Cytokine Signaling (SOCS) family genes. In both RAW264.7 and HMC3, most inflammatory pathway genes were upregulated in cholesterol-treated cells relative to controls, whereas SOCS genes, which act as negative regulators of cytokine signaling, were generally reduced (**Figure 2C-I**). These findings suggest that acute cholesterol not only promotes expression of inflammatory signaling components but may also weaken inhibitory feedback pathways^20^.

We next asked whether these pathway-restricted inflammatory genes were sufficient to distinguish cholesterol-treated from control samples. Principal component analysis (PCA) performed using only the selected inflammatory pathway genes showed clear separation between cholesterol-treated and control cells in both RAW264.7 and HMC3 cells (**Figure 2J-K**), indicating that cholesterol induces a robust and coherent inflammatory transcriptional signature in both cell types.

The most strongly enriched Reactome pathway in both cell lines was “Response of EIF2AK1 (HRI) To Heme Deficiency” (**Figure 2L**). EIF2AK1 is an inflammatory pathway linked to AD^19^. Inspection of the genes contributing to this enrichment showed that the overlapping genes were canonical stress-response genes, including PPP1R15A, DDIT3, TRIB3, CHAC1, ATF3, and ATF4, pointing to activation of integrated stress-response programs. This observation suggested that cholesterol may promote inflammatory activation through induction of cellular stress pathways.

To test this possibility, we organized stress-associated genes into six stress-related subpathways: PERK/ISR/ATF4-CHOP, Heat Shock/Proteostasis, IRE1/XBP1, Oxidative/TXNIP, Stress MAPK/AP-1, and Inflammasome Link. Multiple stress modules were elevated in both RAW264.7 and HMC3, with particularly strong induction of PERK/ISR/ATF4-CHOP and Stress MAPK/AP-1 genes (**Figure 2M**). In several cases, most or all genes within a stress sub pathway were upregulated by cholesterol treatment, indicating coordinated activation of stress-response programs.

Finally, volcano-plot analysis showed that genes linking stress signaling to inflammatory activation were consistently shifted toward positive fold change in both cell types, and several were among the strongest DEGs in the dataset (**Figure 2N-O**). In contrast, treatment with vehicle (MβCD) alone yielded very few differentially expressed genes, and the magnitude and statistical significance of these changes were substantially lower than those induced by cholesterol loading (**Figure S4A-D**). Together, these transcriptional data support a model in which cholesterol induces inflammatory signaling in microglia- and macrophage-like cells, at least in part, through activation of stress-response pathways (**Figure 2I,P**).

### Knockout of astrocyte-derived cholesterol in an AD mouse model

Many primary inflammatory proteins, including PERK, are downstream of cellular cholesterol uptake^12,20,21^. If cholesterol plays a role in neuroinflammation *in vivo*, then knocking out astrocyte-derived cholesterol should reduce microglial activation in an AD mouse model. To test this hypothesis, we examined microgliosis in a familial transgenic (3xTg) mouse model of AD^22^ crossed with our SREBP2^flox/flox^ GFAP-Cre mice^23^. The cholesterol-depleted model uses the human GFAP promoter to drive the expression of Cre recombinase in astrocytes to delete SREBP2 (SB2), which we developed to decrease astrocyte cholesterol synthesis in the brain ^24^.

To confirm knockdown of cholesterol synthesis in primary astrocytes, we visualized astrocytes using GFAP labelling and measured the critical cholesterol synthesis enzyme FDFT1. **Figure 3A** shows FDFT1, a cholesterol synthesis enzyme regulated by SB2, was reduced in SB2−/− astrocytes, confirming the knockdown of the cholesterol synthesis pathway. We analyzed the volume and morphology of astrocyte cells. 3D reconstructions of the hippocampal astrocytes demonstrated a significant 50% decrease in SB2−/− astrocyte volume and number of Sholl intersections compared to wild type (**Figure 3B**). The decrease in volume and intersection correlated with a ~30% decrease in tissue cholesterol (**Figure 3C**).

**Figure 3.**
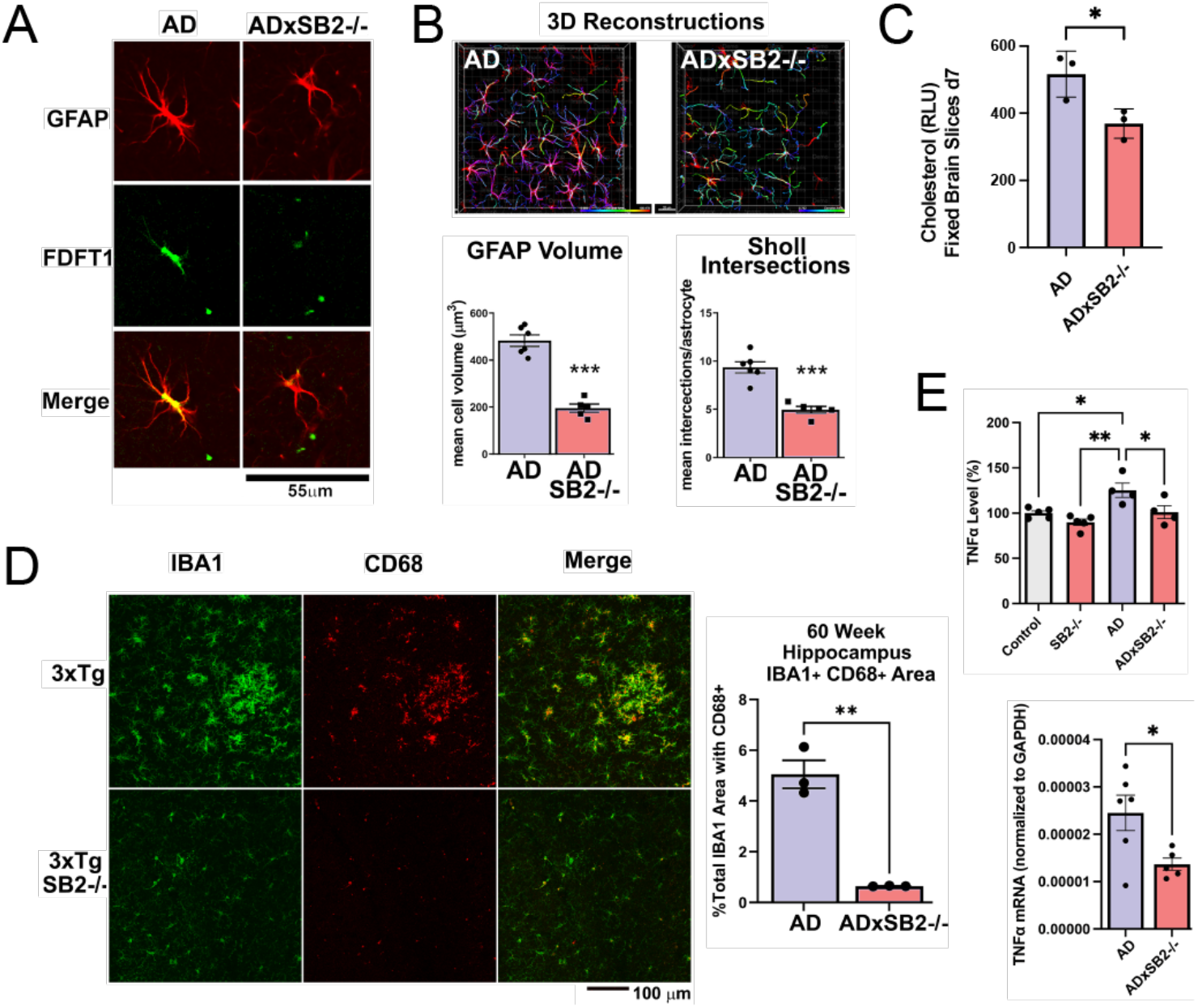
Genetic depletion of SREBP2 in astrocytes inhibits microglial activation in an FAD mouse model. Hippocampal astrocytes and microglia were assessed in 3xTg-AD mice (AD) and AD mice crossed to astrocyte cholesterol deficient Sterol Regulatory Element-Binding Protein (SREBP2)^fl/fl^ GFAP-Cre^+/−^ mice to generate AD x SB2^−/−^. (A) Confocal imaging of astrocytes labelled with GFAP (red) and FDFT1 (green) demonstrates down-regulation of squalene synthase, a key enzyme in the cholesterol synthesis pathway, in AD x SB2^−/−^ astrocytes. (B) Representative images of astrocytes showing a change in morphology and spatial distribution. Analysis of cell volume shows a decrease in astrocyte volume after cholesterol knockdown. Quantitation of astrocyte branches (bar graphs) shows a decrease in branch points, branch length and Sholl intersections in AD x SB2−/− brains compared to AD brains. (C) Fluorescent cholesterol oxidase assay from fixed brain slices. SB2−/− decreases cholesterol compared to AD control. (D) Confocal imaging on brain slices of AD x SB2−/− shows a robust decrease in % CD68+ (a marker of activated microglia) IBA1+ (a marker for total microglia) in SB2−/− animals, suggesting a decrease in activated microglia and macrophages after cholesterol knockdown. (E) (top)TNFα ELISA on brain tissue shows an increased inflammatory level in AD brains compared to wildtype brains. SB2 knockout in astrocytes of AD (ADxSB2−/−) brains significantly downregulated TNFα. (bottom) qPCR of inflammatory cytokine TNFα. Comparison made with a Student’s T test; or ANOVA (E, top) *p≤0.05, **p≤0.01, ****p≤0.0001.

We next immunostained these brains for IBA1 and CD68 proteins to assess activation of the hippocampal microglia and infiltrating macrophages. IBA1 is a marker of total microglia and CD68 is a marker of phagocytosing microglia ^25,26,27^. Imaging showed a robust decrease (~5 fold) in CD68 for the ADxSB2−/− brains (**Figure 3D**), suggesting a knockdown in astrocyte cholesterol decreases the activation of microglia and macrophages.

To confirm knockout of the astrocyte-derived cholesterol reduced the release of inflammatory cytokines *in vivo*, we measured the level of TNFα. Using ELISA, we observed that with WT astrocyte-derived cholesterol, the AD brains had significantly higher TNFα levels compared to controls (**Figure 3E**, top). Without astrocyte cholesterol, TNFα levels were reduced to the level of control brains (**Figure 3E**, top). Consistently, we found TNFα mRNA levels were also significantly lower in ADxSB2−/− brains compared to AD brains (**Figure 3E**, bottom). Other cytokines, including IL1β, IL1α, INFα and INFβ, showed the same trend in mRNA levels, although the changes did not reach statistical significance (**Figure S6A-D**).

### Knockout of astrocyte-derived cholesterol during acute systemic inflammation

Acute inflammation in the periphery is known to activate microglia in the brain causing neuroimpairment^28^. This is thought to include opening of the blood brain barrier (BBB) and infiltration of immune cells^1,2^. Cholesterol has also been speculated to cross the BBB in these situations and increase neurodegeneration^12,29^ neuroinflammation, we injected control (SREBP2flox) and SB2−/− mice intraperitoneally (i.p.) with a single 2 mg/kg dose of LPS and examined microglia activation in the brains of these animals after 24 hours and 7 days (**Figure 4A**). To quantitate microglia activation, we measured CD68 levels in IBA1+ cells by confocal immunofluorescence as described above.

**Figure 4.**
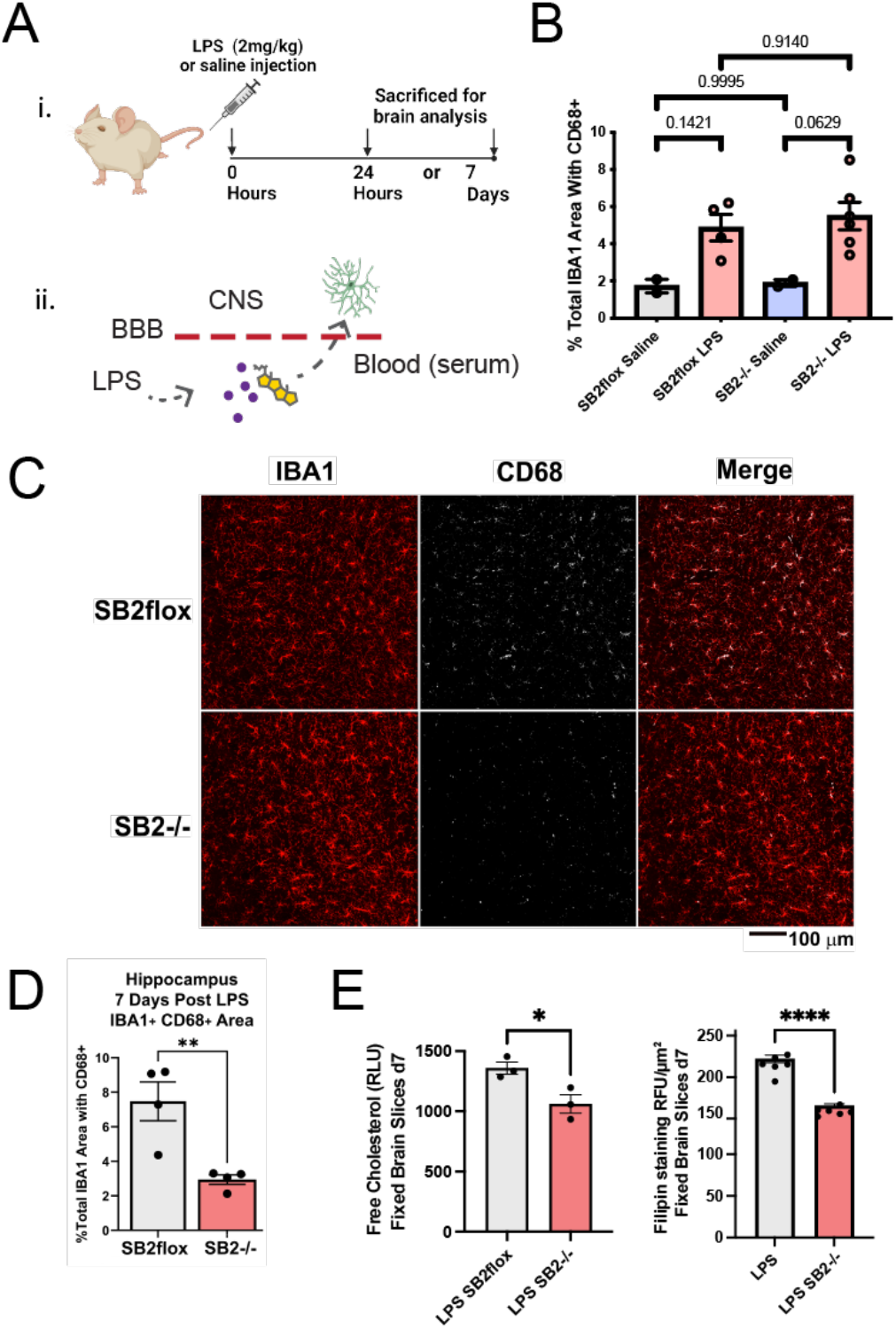
Decreased astrocyte cholesterol helps facilitate recovery from neuroinflammation after peripheral LPS injection. SREBP2 floxed mice (SB2flox) were crossed to GFAP-Cre^+/−^ mice to generate mice that do not produce cholesterol specifically in astrocytes (SB2−/−). (A) i. timeline of 2 mg/ml LPS treatment intraperitoneal (I.P.). Hippocampal microglia were assessed for CD68 and IBA1 content after 24 hours and 7 days. ii. Model of BBB opening in response to LPS allowing passage of serum and immune cells. (B) Quantification of CD68 staining in IBA1 positive cells from confocal images (see Supplemental Figure S6B) of SB2−/− mice 24 hours after LPS treatment. (C) Confocal images day 7. (D) Quantitation of CD68 staining in IBA1 labeled cells is decreased 3-fold in SB2−/− mouse brains by day 7. (E) Fluorescent cholesterol oxidase assay (left) and filipin staining (right) from fixed brain slices. SB2−/− decreases cholesterol in LPS-treated animals. Each point is a biological replicate (n=3-6 animals per experiment).

24 hours after injection with LPS we observed a large increase in CD68 positive IBA1 cells, but no difference between SB2flox control and SB2−/− mice (**Figure 4B, S7A-B**), suggesting peripheral cholesterol and cytokines entered the brain following LPS injection and astrocyte-derived cholesterol is not necessary for microglia activation. In contrast, after 7 days, LPS-injected SB2−/− mice showed significantly less CD68 in IBA1+ cells compared to controls (**Figure 4C-D**). Both a cholesterol oxidase assay and filipin staining showed a similar decrease in brain cholesterol in LPS-injected SB2−/− compared to LPS-injected controls (**Figure 4E**). This strong correlation of cholesterol both with and without activation further supports cholesterol’s role in microglial activation and suggests that decreasing astrocyte-derived cholesterol may help resolve neuroinflammation and the neurotoxic effects of reactive astrocytes.

Of note, there is no difference in apoE in the cerebrospinal fluid (CSF) between SB2flox controls and SB2−/− mice, thus the difference in response across genotypes cannot be explained by a change in apoE levels (**Figure S7E-F**).

### Uptake of astrocyte-derived cholesterol into immune cells

Next, we confirmed the cholesterol released from the activated astrocytes is taken up into cultured immune cells. First, we activated cultured astrocytes with a cytokine cocktail (IL1β, TNFα, and IL6), then conditioned fresh media with the activated astrocytes, and finally applied the reactive ACM (rACM) to macrophages and measured cholesterol uptake into the cultured cells.

In RAW264.7 (RAW) the application of rACM produced a robust uptake of cholesterol compared to control ACM with no prior exposure to cytokines. Cellular cholesterol levels increased almost 30% (**Figure 5A**). In microglia, the same rACM induced only a small (~2%), albeit statistically significant, increase in cellular cholesterol levels (**Figure 5B**). This small percentage of cholesterol uptake from rACM into HMC3 was not due to a lack of available cholesterol in the rACM evident by the uptake into RAW cells, suggesting HMC3 require additional input or distinct conditions for taking up cholesterol. Direct application of cytokines to macrophages and microglia had no effect on cellular cholesterol levels (**Figure S2E-F**), hence the increases in cellular cholesterol with rACM treatment is likely exogenous cholesterol release from the astrocyte and not de novo synthesis of cholesterol in macrophages or from cytokines^1,2^.

**Figure 5.**
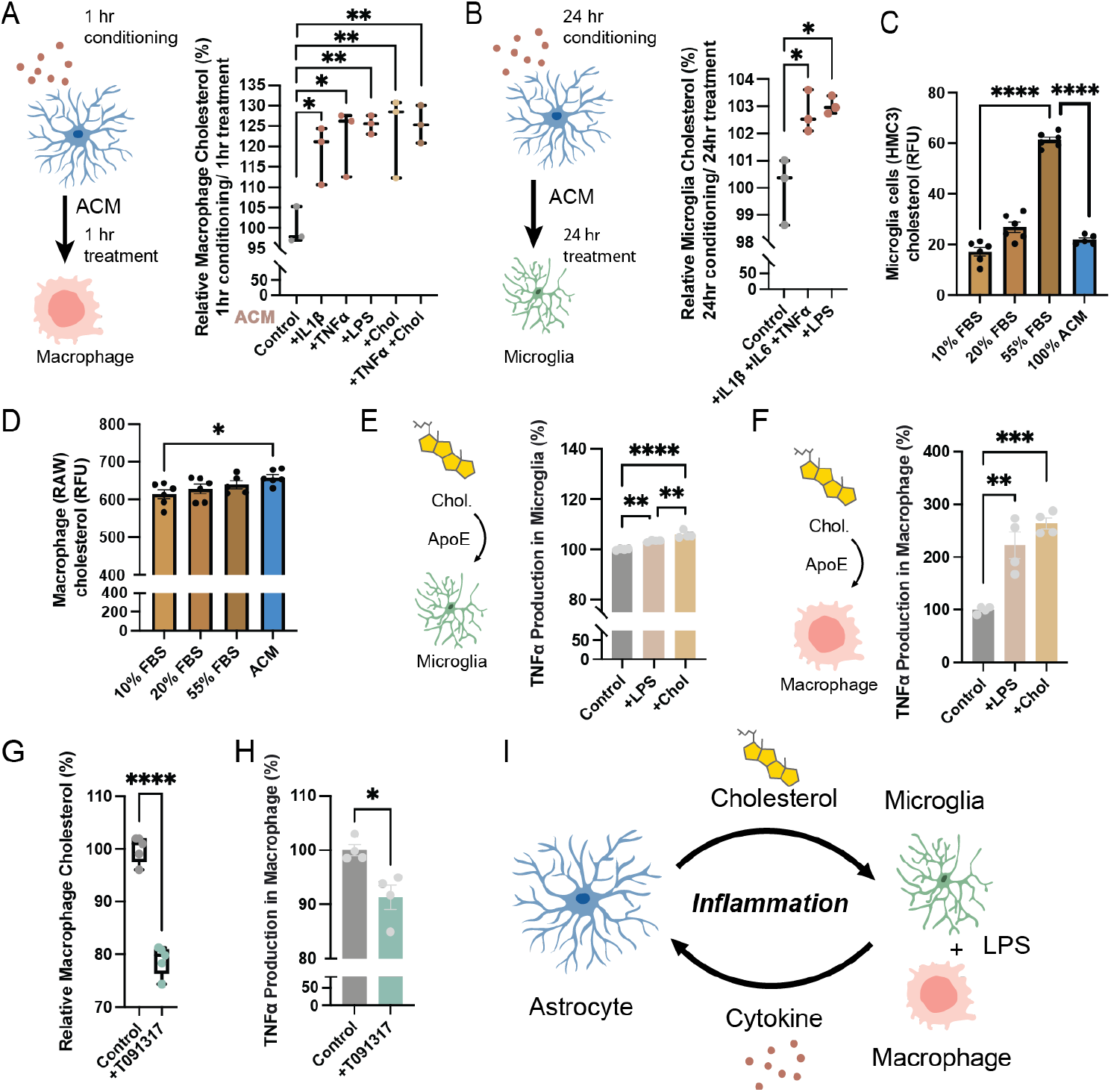
Reactive astrocytes promote uptake of cholesterol in macrophages. (A) Experimental design and cholesterol uptake assay in RAW264.7 macrophages. Primary astrocytes (blue shaded cell) were stimulated with cytokines (100 ng/mL, brown dots) for 1 h, washed, and incubated in fresh media for an additional 1 h to generate reactive astrocyte-conditioned media (rACM). The rACM was transferred to RAW264.7 macrophages (pink shaded cell) for 1 h, and cellular cholesterol was measured using a fluorescent cholesterol oxidase assay. (B) Experimental design and cholesterol uptake assay in HMC3 microglia. Primary astrocytes were stimulated with cytokines (100 ng/mL) for 24 h, washed, and incubated in fresh media for an additional 24 h to generate rACM. The rACM was transferred to HMC3 cells (green shaded cell) for 24 h, and cellular cholesterol was measured using a fluorescent cholesterol oxidase assay. (C-D) Free cholesterol uptake into (C) HMC3 cells and (D) RAW264.7 macrophages measured by fluorescent cholesterol oxidase assay. Blood is ~55% serum. (E-F) Measurement of TNFα protein release after direct treatment of microglia (E) and macrophages (F) with cholesterol (4 ng/ml apoE + 10% serum). Cholesterol behaved similar to 10 ng/mL LPS (positive control) in both cell types. (G-H) T091317, a Liver X receptor agonist, decreased cholesterol in macrophages, which correlated with decreased TNFα production by macrophages. (I) Proposed model for astrocyte cholesterol in neuroinflammation. Astrocyte-derived cholesterol induces activation of microglia and macrophages and triggers cytokine release. Cytokines and other pro-inflammatory molecules released by microglia and macrophages in turn stimulate cholesterol release from astrocytes, triggering neuroinflammation. Values are expressed as mean. In A-B, each point is a well of cells from 2 animals done at least in duplicate (>4 animals). Comparison made with a Student’s T test; *p≤0.05, **p≤0.01, ***p≤0.001, ****p≤0.0001.

### Uptake of serum cholesterol into immune cells

As mentioned, AD is associated with a leaky blood brain barrier (BBB) and cholesterol from the periphery is thought to contribute to the progression of the disease^1,2,12^. Using the cholesterol oxidase enzyme assay, we tested the ability HMC3 microglia to uptake cholesterol from fetal bovine serum (FBS), a source of peripheral cholesterol. In mammalian blood, serum makes up ~55% of the total volume.

We found, cholesterol uptake into microglia increased at 20% and dramatically at 55% serum containing media, suggesting under conditions where HMC3 are exposed to serum, they can take up significant amounts of cholesterol (**Figure 5C**). At low serum (10%), the concentration typically used in culture systems, HMC3 cells took up approximately the same cholesterol compared to incubation with ACM. Interestingly cultured RAW 264.7 macrophage did not respond appreciably to serum over ACM—even at the highest concentration, serum (55%) uptake was less than ACM (**Figure 5D**).

### Cholesterol activation of microglia and macrophages

To confirm microglia activation by apoE-mediated cholesterol uptake, we cultured HMC3 cells with apoE3 with/without FBS and measured TNFα production using an enzyme-linked immunosorbent assay (ELISA) for TNFα. As mentioned, we previously showed that cholesterol requires apoE for uptake into neuronal membranes^23^, making it a reasonable test for physiological uptake of cholesterol into immune cells here.

We found that treatment of microglia with apoE and 10% FBS significantly increased TNFα production (**Figure 5E**). As a control we added LPS, which activates TNFα production in cultured microglia ^30^. The effect of apoE and FBS was stronger than LPS incubation alone (**Figure 5E**), a result consistent with our observation that cholesterol and inflammation combine to activate microglia. The purified apoE3 had <0.1ng/µg endotoxin (LPS) (per manufacturer testing) which is <0.0004 ng/mL final concentration and insignificant compared to the 10 ng/mL LPS we added in the assay.

To directly compare the cholesterol response of microglia with macrophages, we incubated RAW 264.7 macrophages with apoE and cholesterol. We found that the uptake of cholesterol into macrophages with apoE was dramatic, >200% (**Figure 5F**). The generation of secreted TNFα using purified apoE3 was remarkably consistent with the relative increases in cholesterol seen using rACM from activated astrocytes (**Figure 5A-B**). These results further suggest the amount of cytokine production correlates to the amount of cholesterol uptake by the cell.

In addition to uptake, efflux can regulate the cellular levels of cholesterol. We tested the effect of cholesterol efflux on immune cells function. Efflux of cholesterol in microglia can be induced with a liver X agonist. Liver X protein upregulates the ATP binding cassette (ABC) subfamily G member 1 (ABCG1) transporter among many other genes, so the assay is not specific^37^. Nonetheless, in RAW macrophages ABC subfamily A member 1 (ABCA1) is a major contributor to cholesterol efflux, a function that should affect TNFα levels.

To test cholesterol efflux, we treated RAW macrophages with the Liver X receptor agonist T0901317 and measured cholesterol levels using the fluorescent cholesterol oxidase assay. Consistent with activation of ABCA1, we saw a 20% decrease in macrophage cholesterol (**Figure 5G**) which corresponded to a ~10% decrease in TNFα levels (**Figure 5H**). This again supports the hypothesis that cytokine production is regulated by the levels of cholesterol in immune cells (**Figure 5I**).

While astrocytes are the primary source of cholesterol in the brain, and we saw no increases in microglial cholesterol with cytokine treatment (**Figure S2D**), microglia do possess the machinery for cholesterol synthesis. To determine if microglia cholesterol synthesis could contribute to TNFα secretion we treated SIMA-9 cells, a mouse microglia-derived cell line, with zaragozic acid to inhibit cholesterol synthesis. Cells in serum containing media were then treated with LPS and the conditioned media was assayed for TNFα. We observed effective blockade of cholesterol synthesis as evidenced by compensatory upregulation of FDFT1, a key enzyme in the cholesterol synthesis pathway (**Figure S7C**). SIMA-9 cells produced a robust increase in TNFα release into the media in response to LPS, which was unchanged by pretreatment with zaragozic acid (**Figure S7D**), demonstrating a reliance on uptake of extracellular cholesterol over endogenous cholesterol synthesis for the secretion of TNFα and consistent with the results in **Figure 5E-F**.

### Cholesterol amplifies interferon signaling through nanoscale organization of interferon receptors

Having shown that cholesterol induces cytokine pathways in immune cells and that inflammatory cytokines induce cholesterol synthesis in astrocytes, we next sought to investigate the genomic pathways linking cholesterol metabolism and inflammatory responses. To manipulate intracellular cholesterol levels, we inhibited cholesterol synthesis with simvastatin, an inhibitor of HMG-CoA reductase, a key step in the cholesterol synthesis pathway, or increased intracellular cholesterol through genetic deletion of ABCA1, a major cholesterol efflux transporter^4,31^.

Proteomics analysis of U251MG human astrocyte-like cells revealed a strong relationship between cholesterol metabolism and interferon signaling (**Figure 6A-B**). Inhibition of cholesterol synthesis with simvastatin resulted in a broad reduction in interferon-stimulated gene (ISG) protein levels. Conversely, increasing intracellular cholesterol through ABCA1 knockout increased the abundance of many ISGs. These reciprocal effects indicate that intracellular cholesterol levels strongly regulate interferon-responsive transcriptional programs and identify interferon signaling as a cholesterol-sensitive inflammatory pathway. This is consistent with our transcriptomic analysis of cholesterol-loaded HMC3 microglia, which showed increased expression of key interferon pathway genes, including interferon regulatory factors (IRFs) and interferon-stimulated genes (ISGs) (**Figure 2F-G**).

**Figure 6.**
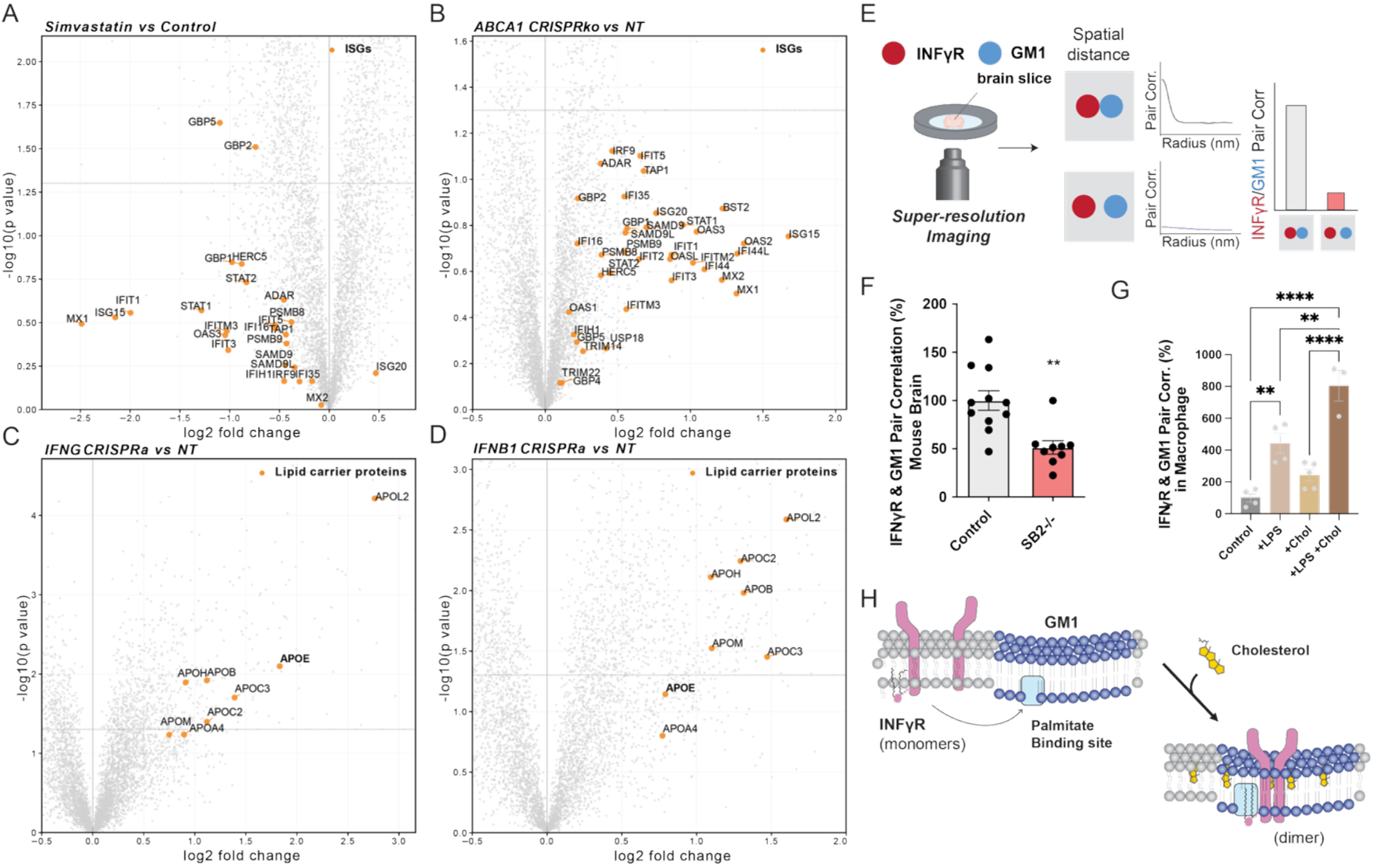
Cholesterol enhances interferon signaling through nanoscale organization of IFNγR. (A) Volcano plot showing differential protein abundance in U251MG human astrocyte-like cells treated with simvastatin compared to untreated controls. Interferon-stimulated genes (ISGs) are highlighted in orange and labeled. Inhibition of cholesterol synthesis resulted in broad downregulation of ISG proteins, indicating suppression of interferon-responsive signaling pathways. (B) Volcano plot showing differential protein abundance in ABCA1 knockout U251MG cells compared to non-targeting (NT) controls. ISGs are highlighted in orange and labeled. Genetic disruption of cholesterol efflux increased the abundance of numerous ISGs, demonstrating that intracellular cholesterol accumulation promotes interferon-responsive transcriptional programs. (C) Volcano plot showing differential protein abundance following CRISPR activation (CRISPRa) of IFNG in U251MG cells relative to NT controls. Apolipoproteins are highlighted in orange and labeled. Activation of IFNG increased expression of multiple apolipoproteins, including apoE, consistent with enhanced cholesterol transport capacity. (D) Volcano plot showing differential protein abundance following CRISPRa of IFNB1 in U251MG cells relative to NT controls. Apolipoproteins are highlighted in orange and labeled. Similar to IFNG activation, IFNB1 activation increased expression of multiple apolipoproteins, including apoE. (E) Experimental setup and analysis for two color dSTORM. GM1 lipid and protein were fluorescently labeled and their localizations determined by dSTORM. High correlation at short distances (5-10 nm) is graphed and indicates co-localization. (F) dSTORM imaging of SB-flox (Control) and SREBP2 KO (SB2−/−) mouse brain tissue demonstrates reduced IFNγR-GM1 clustering in SB2−/− brains. (G) dSTORM imaging of cultured immune cells in 8-well chambers. LPS treatment or cholesterol loading alone shows a trend in increasing IFNγR-GM1 lipid cluster association. LPS and cholesterol together has an additive effect. (H) The GM1 lipid raft is the platform to initiate inflammatory signaling. (F) each point is a cell from 2 experimental brains (4 animals), done in duplicate. (G) each point is a biological replicate (single cell) and comparison are made from samples prepared at the same time. Each experiment was done at least in duplicate. Statistical comparison made with a Student’s T test or ANOVA (F,G); *p≤0.05, **p≤0.01, ***p≤0.001, ****p≤0.0001.

Because of apoE’s role in cholesterol transport, we next tested whether interferon signaling reciprocally regulates apolipoprotein expression. CRISPR activation of either IFNG or IFNB1 in U251MG cells increased the abundance of all apolipoproteins detected by mass spectrometry, including apoE (**Figure 6C-D**). Together, these data support a bidirectional relationship in which cholesterol promotes interferon-responsive signaling, while interferon signaling enhances cholesterol transport capacity.

We next investigated how cholesterol enhances interferon signaling at the molecular level. Cholesterol and saturated lipids promote the formation of ordered regulatory nanodomains, often referred to as lipid rafts, which facilitate receptor clustering and signal transduction and thereby enhance inflammatory signaling pathways^12^ (**Figure S1C**). Several inflammatory receptors, including TLR4, TREM2, and IFNγR, exhibit cholesterol-dependent membrane regulation ^32^. IFNγR is palmitoylated and activated in lipid rafts by dimerization. Clustering activates IFNγR and TLR4 independent of their ligands by dimerization, while clustering activates TNFα and TREM2 by exposing them to their hydrolytic enzymes^33^ (**Figure S8A, S1D**).

We hypothesized that astrocyte-derived cholesterol regulates interferon signaling by controlling the localization of IFNγR within lipid rafts, a mechanism analogous to the cholesterol-dependent trafficking of amyloid precursor protein (APP) in neurons^15,37,38^. To analyze cholesterol-induced changes in spatial location of IFNyR, we used direct stochastic optical reconstruction microscopy (dSTORM), a super-resolution imaging approach that allows ample sub-diffraction limited observations suitable for observing the nano-environment of proteins and detecting nanoscopic movements (<200 nm) between cholesterol dependent GM1 and cholesterol independent PIP_2_ containing lipid regulatory domains^34^. We recently employed similar dSTORM techniques to establish membrane-mediated mechanisms of amyloid production, anesthesia, and mechanosensation ^17,23,35,36^.

Using dSTORM super-resolution imaging and cross pair correlation analysis (**Figure 6E**), we found that SB2−\− reduced IFNγR association with GM1-rich lipid rafts by 50% *in vivo* (**Figure 6F**). The localizations were determined from aged (60-week-old) wild-type and astrocyte-specific SREBP2 knockout (SB2−/−) mice. In wild-type tissue, IFNγR showed measurable association with GM1-positive lipid domains, whereas astrocyte-specific depletion of cholesterol significantly reduced IFNγR-GM1 pair correlation. These findings demonstrate that astrocyte-derived cholesterol contributes to the nanoscale membrane organization of IFNγR *in vivo*.

To determine whether cholesterol promotes IFNγR recruitment into lipid rafts under inflammatory conditions, we treated RAW264.7 macrophages with apoE-mediated cholesterol loading and/or LPS stimulation and quantified IFNγR colocalization with GM1 lipids using dSTORM imaging. Cholesterol loading or LPS treatment alone increased IFNγR association with GM1 domains, while the combination produced the strongest effect (**Figure 6G**). These findings suggest that maximal IFNγR recruitment into cholesterol-rich membrane domains requires both inflammatory signaling and elevated cholesterol availability.

Together, these data support a model in which astrocyte-derived cholesterol amplifies interferon signaling by promoting nanoscale reorganization of IFNγR into cholesterol-rich membrane domains (**Figure 6H**).

### Clustering of inflammatory proteins in an FAD mouse model

Lastly, we asked whether cholesterol-dependent membrane reorganization extends to inflammatory mediators TREM2 and TLR4 in an FAD mouse model. Wt or FAD mice were crossed with astrocyte-specific SREBP2 knockout mice (ADxSB2−/−)(same tissue as in **Figure 3**). At 60 weeks of age, brain sections (40 µm) were prepared for dSTORM imaging and compared with age-matched wild-type and FAD controls.

We included two types of controls; SB2-flox (Control), with intact SREBP2 and no FAD mutations, this comparison shows the ability of astrocyte-derived cholesterol to regulate the proteins in a non-disease state, and two, an FAD alone control (AD, intact SREBP2); which reveals cholesterol’s ability to ameliorate the FAD state when compared to the astrocyte SREBP2 −/− crossed to the FAD (ADxSB2−/−).

In a diseased state with normal cholesterol (ADxSB2-flox) TREM2 showed increased association with GM1 lipid clusters (**Figure 7A**). Astrocyte-specific deletion of cholesterol (ADxSB2−/−) significantly decreased TREM2’s association with GM1, demonstrating at the molecular level that astrocyte-derived cholesterol regulates the nanoscale localization of TREM2 *in vivo*. Compared to SB2-flox control TREM2\GM1 pair correlation was similar in SB−/− mice suggesting the association of TREM2 with GM1 domains was already low in wild-type tissue.

**Figure 7.**
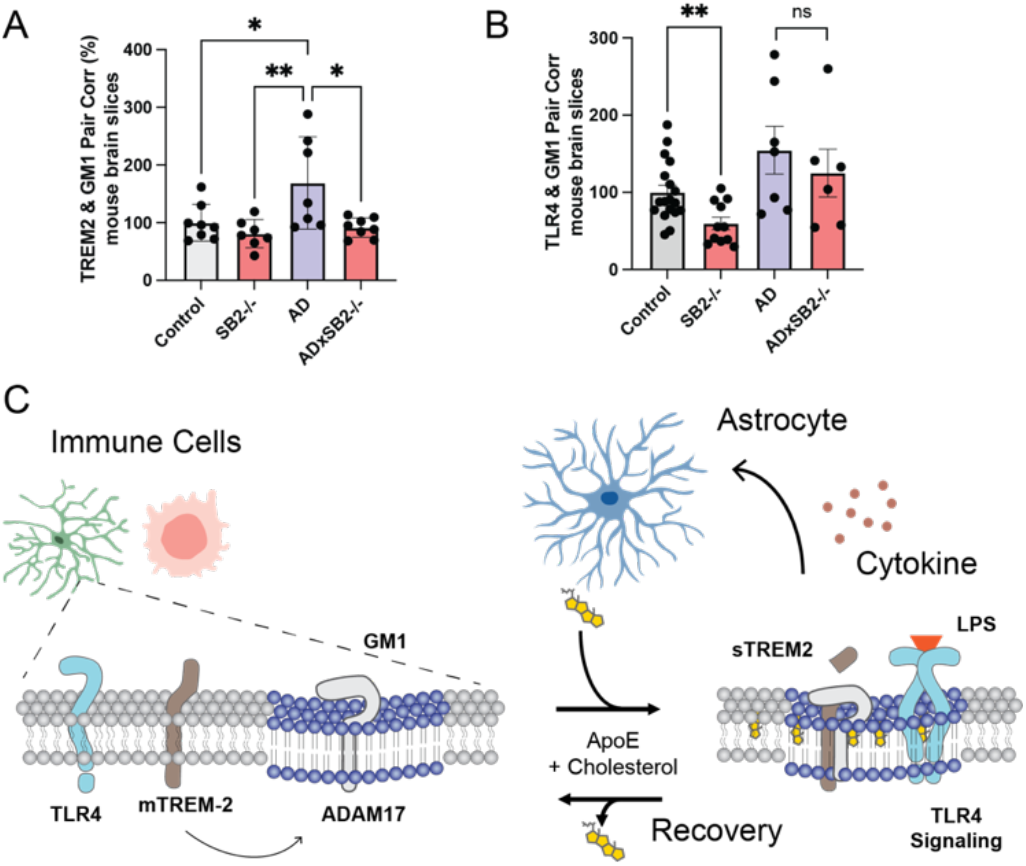
Cholesterol’s spatial regulation of inflammatory proteins in AD mouse brain. (A-B) dSTORM super resolution imaging on mouse brain slices. (A) Compared to control animals (white shading), clustering of TREM2 with GM1 regulatory domains significantly increased in 3xTG (AD, purple shading). In AD animals with astrocyte SREBP2 (cholesterol synthesis) knockout (AD x SB2−/−) (red shading) the clustering of TREM2 was reversed. (B) SB2−/− affects TLR4 primarily in control tissue but not significantly in the AD mouse model. (C) Model depicting GM1 domain regulation by astrocyte-derived cholesterol. At nanoscopic levels, TLR4 and membrane bound TREM2 (mTREM2) sort into lipid compartments away from ADAM17. ADAM17 is a protease that hydrolyzes mTREM2. Upon inflammatory stimulation (e.g., lipopolysaccharide (LPS)) immune cells in the brain release cytokines which make their way to astrocytes and induce release of cholesterol. The cholesterol from astrocytes, activates microglia causing a clustering of TLR4 to allow cytokine binding and mTREM2 with its hydrolytic enzyme ADAM17, producing more cytokines.

In contrast TLR4’s association with GM1 lipids in control (SB2-flox) was almost 2-fold higher compared to SB−/−, suggesting TLR4 has significant association with GM1 lipid in non-FAD mice (**Figure 7B**). The TLR4/GM1 pair correlation in AD mice did appear to increase compared control, but did not reach statistical significance, likely because the control already has significant pair correlation. However, TLR4/GM1 pair correlation in ADxSB2−/− only decreased slightly compared to AD mice, indicating TLR4 likely has inputs other than cholesterol in the diseased state.

To confirm cholesterol’s molecular model in a controlled cellular environment, we treated cultured HMC3 microglia and RAW264.7 macrophages with apoE-mediated cholesterol loading and measured colocalization of TREM2 with GM1 lipids (**Figure S8B-C**). In both HMC3 microglia and RAW264.7 macrophages, cholesterol loading or LPS stimulation increased TREM2 association with GM1 domains, while the combination produced the strongest effect (**Figure S8B-C**). These data from cultured cells suggest that lipidated apoE contributes to the nanoscale clustering of inflammatory receptors.

## Discussion

The data presented here establish cholesterol as a secreted regulatory factor integral to the escalation and maintenance of neuroinflammation. **Figure 5I** shows a combined cellular molecular model based on the data presented here and the known function of astrocytes, microglia, and macrophages. Cytokines from microglia and macrophages activate the synthesis and release of astrocyte-derived cholesterol (**Figure 1A-B**). Cholesterol uptake into immune cells, while nuanced by conditions and cell type, enhances cytokine release suggesting a positive feedback loop that results in the escalation of neuroinflammation.

Abnormal lipid accumulation in neurodegeneration was first observed more than 100 years ago by Alois Alzheimer ^37^. Alzheimer described an extensive accumulation of ‘adipose saccules’ in the patient’s brain, which is a major store of cholesterol and fatty acids ^38^. The studies here give a plausible role and mechanism of action for the elevated lipids.

The uptake of astrocyte-derived cholesterol into immune cells resembles that of neuronal uptake and induced amyloid production by neurons (**Figure S8A**) ^15^. Secretions from activated astrocytes also include saturated fats which may contribute to the molecular mechanism proposed in (**Figure 7C**). The signaling role of cholesterol established here and elsewhere likely serves as a link between amyloid and inflammatory pathologies. ^42,43^

Our data suggest that in addition to brain-derived cholesterol, peripheral cholesterol may play an important role in perpetuating neuroinflammatory responses in disease. The uptake of serum cholesterol into cultured microglia (**Figure 5C**) and ACM in macrophages (**Figure 5D**) were distinct. The activated microglia in SB2−/− KO after LPS injection agree with peripheral cholesterol promoting AD through an open BBB, as previously speculated^12,29^. But brain cholesterol clearly plays a role, e.g., the reduced microglia activation at day 7 after LPS injection in SB2−/− mice (**Figure 4C-D**) which only lack astrocyte cholesterol. Hence in the ADxSB2−/− mouse model the lack of inflammation is likely due to a decrease in cholesterol dependent inflammatory signaling.

The molecular mechanisms driven by cholesterol are still emerging. We have tested only a few palmitoylated cytokines and determined their nano-environment and regulation (**Figure 7, S1E**). Many other cytokines are palmitoylated and regulated by lipid rafts (see supplemental discussion), similar to TNFα. Presumably cholesterol affects many proteins at the same time using this palmitate-mediated clustering mechanism, e.g. ion channel, transporters, and G-proteins.

Future studies will be needed to test cholesterol’s overall role in driving hyperexcitability through altered metabolic state in nerves. And *in vivo* studies will be needed to better characterize the source of inflammation entering the brain from the periphery and the role of microglia and macrophages in responding to BBB opening^44,45^.

## Acknowledgements

We thank Julian Bois for help with the cholesterol assays and Yutao Wang and Allan Tall for reading of the manuscript and helpful suggestions. We thank Tajie Harris for assistance with CSF experiments. This work was supported by the NIH via an R21 (Grant R21 AG078845-01), Director’s New Innovator Award (DP2NS087943), and R01 (Grant R01NS112534) to S.B.H., a K08 (Grant K08DK097293), R01 (Grant R01AG080773) and Owens Family Foundation Award to H.A.F., and a NIH T32 (Grant T32DK764627) to J.A.K. We are grateful to the JPB Foundation for the purchase of a super resolution microscope.

## Supplementary Information

### Discussion

The dSTORM used here is an important technical advancement. First, we measured a cholesterol dependent inflammatory state using the clustering of inflammatory receptors. The signal is direct from the tissue and at the source. Most inflammation is measured by downstream effectors such as RNA expression, but these are complicated by a myriad of downstream inputs that don’t necessarily indicate the original inflammatory state in the tissue of interest. While RNA expression and ELISA are valuable tools, particularly when there is robust inflammation, such as with infection, dSTORM imaging is likely to better reveal early or low-grade chronic inflammation, in time dSTORM may be a preferred method of determining an early inflammatory signal.

Second, we have directly labeled the lipids and compared the protein localization to the signature lipid (GM1). Fluorescent probes targeted to GM1 lipids with palmitate are insufficient as the palmitoylated proteins can leave the GM1 domain when cholesterol conditions change^39^. Hence the ability to function as a GM1 sensor is lost. By directly labeling the lipid, we can determine nanoscopic trafficking in the membrane. It is the nanoscopic movement that appears to control the inflammation.

For our studies the localization of the proteins dictates their function. The location was determined by fixing the cells and measuring pair correlation of the protein with the lipid. The lipids are relatively small, and they are best labeled with antibodies or, in the case of GM1 lipids, with cholera toxin B (CTxB). CTxB is pentadentate which is why the label has very high affinity to a small lipid. The labeling can cause some clustering even when fixed prior to labeling. We have seen very minimal clustering in our setup in multiple cell types ^40^. Nonetheless, artificial clustering does not adversely affect localization of proteins since it has very little effect on where a protein is localized. In fact, in 2003 Reinhard Jahn intentionally used clustering of unfixed cells to help define if proteins were colocalized or not. He called this “antibody patching”. In short, he induced the clustering and then looked to see which proteins clustered with the different “patches” of antibodies ^41^. The patching was necessary since at the time they didn’t have super resolution imaging. It is evident from those studies that even if there is some residual clustering after we fix, that is very unlikely to affect the relative localization of fixed proteins. Since our conclusions are based off pair correlation and localization, not size, any putative clustering by CTxB is likely desirable for determining an accurate result.

## Supplementary Figures

**Figure S1.**
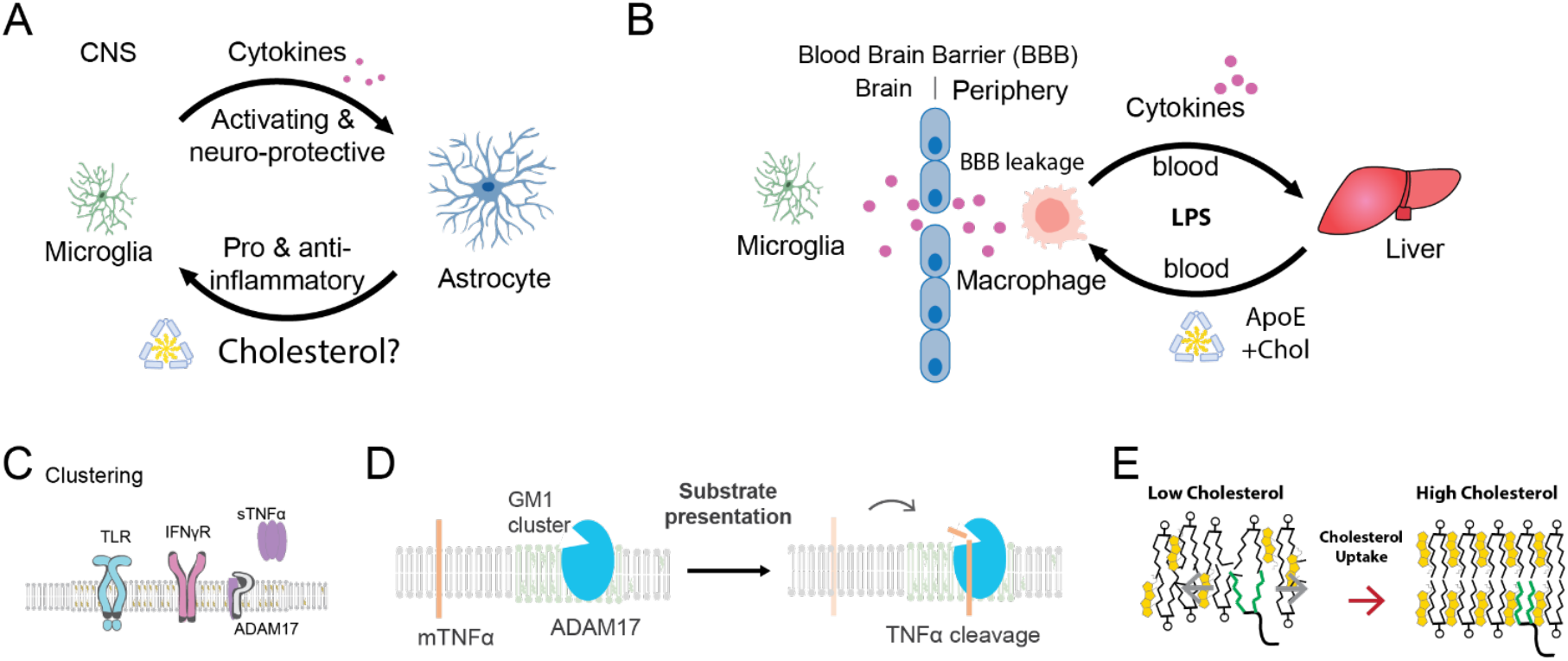
Molecules of neuroinflammation. (A) Communication between microglia and astrocytes. Astrocytes (blue shaded cell) and microglia (green shaded cell) interact by releasing signaling factors locally, known as paracrine signaling. Microglia primarily release cytokines. Astrocytes release both pro and anti-inflammatory factors ^7^. Cholesterol (yellow shading) is drawn in a complex with apolipoprotein E (apoE, light blue shading). The current study asks if astrocytes produce cholesterol that makes its way to microglia and is this a signal to microglia? (B) Inflammation in the CNS is influenced by the periphery. Lipopolysaccharide (LPS) and cytokines induce cholesterol synthesis in the liver. The liver produces cholesterol which is released and transported in the blood in lipoprotein particles containing apoE and other lipoproteins (not shown). ApoE binding to cell surface receptors on macrophages (not shown) and uptake of the cholesterol activates the macrophages in the periphery, producing cytokines. Through poorly understood mechanisms, LPS in the periphery can activate microglia in the CNS. (C) A representation of cytokine receptors regulated by clustering in lipid rafts. Cholesterol induces clustering of inflammatory proteins. In the case of toll-like receptor (TLR) the clustering causes dimers to form and this leads to autophosphorylation. In the case of tumor necrosis factor alpha (TNFα) a membrane tethered TNFα (mTNFα) clusters with its hydrolytic enzyme ADAM17 where the enzyme can catalyze the production of soluble TNF*α* (sTNFα). sTNFα can then signal inflammation. D) Substrate presentation is a biological mechanism that activates a protein by gaining access to its substrate ^42^ (e.g., ADAM17 accessing mTNFα) ^43^(E) Molecular basis for cholesterol’s ability to cluster palmitoylated proteins. Cholesterol orders the acyl chains of saturated lipids. The order is thought to create a low energy state. The order of the lipids creates a binding site for palmitates. The cell covalently attaches palmitates to proteins in a process called palmitoylation and the palmitoylation imbues the protein with an affinity for cholesterol dependent ordered lipids ^44^. Increasing cholesterol in the membrane increases the number of palmitate binding sites and drives proteins to cluster with GM1 lipids.

**Figure S2.**
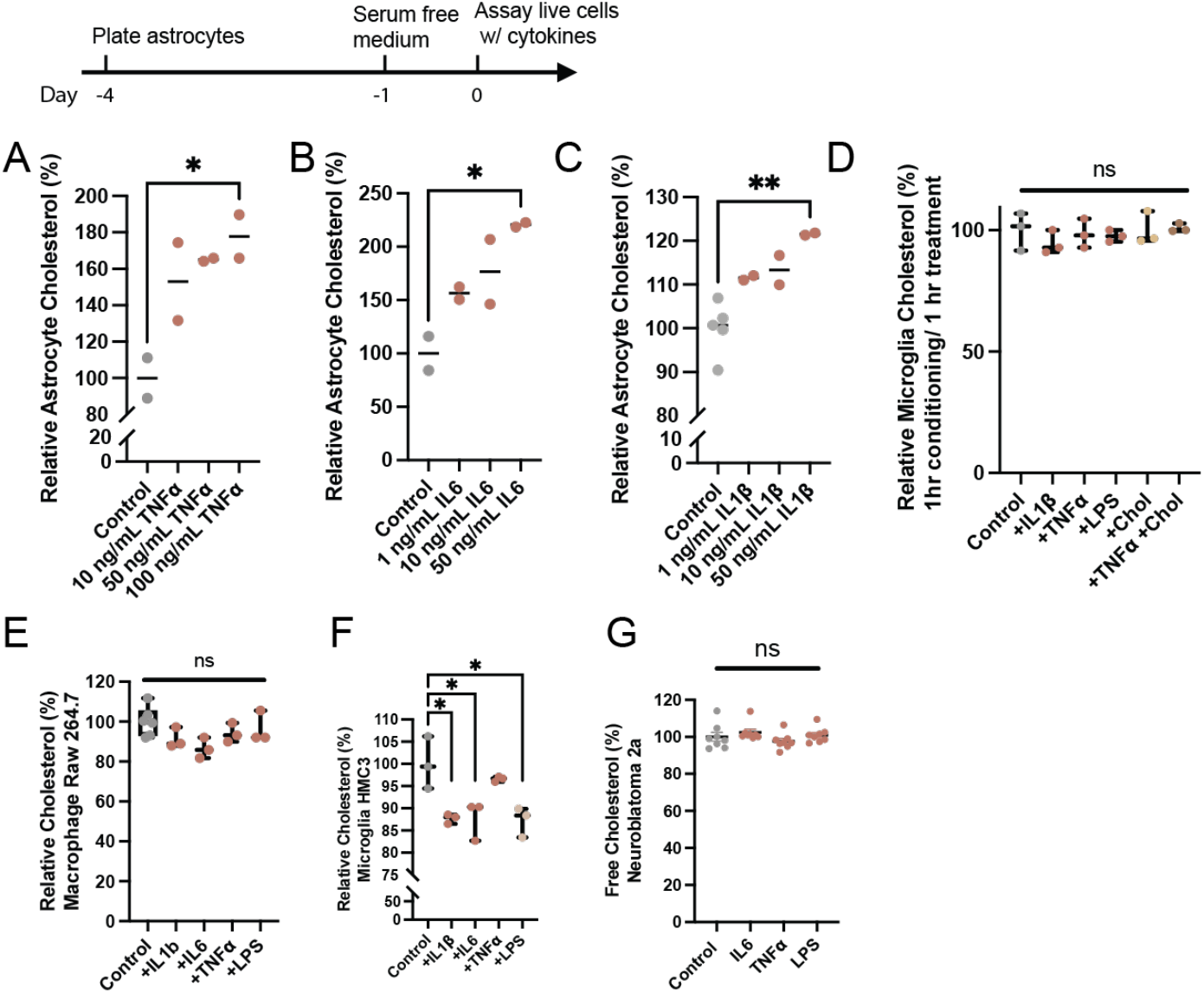
Cytokine dose responses. (A-C) Combined cholesterol from media and available in the plasma membrane measured by a fluorescent cholesterol assay. Pro-inflammatory cytokines IL1β, IL6, and TNFα dose dependently induce cholesterol synthesis in astrocytes. (D) Cytokines (50 nM) were added to primary astrocytes and then the astrocyte-conditioned media (ACM) was placed on HMC3 microglia for 1 hour and then the cellular cholesterol was measured using a fluorescent cholesterol oxidase assay. (E-G) Free cholesterol levels in the cellular membranes of RAW 264.7 macrophages (E) and microglia HMC3 (F), and Neuroblastoma 2a (N2a)(G) after direct application of 100 ng/mL cytokines in serum free media.

**Figure S3.**
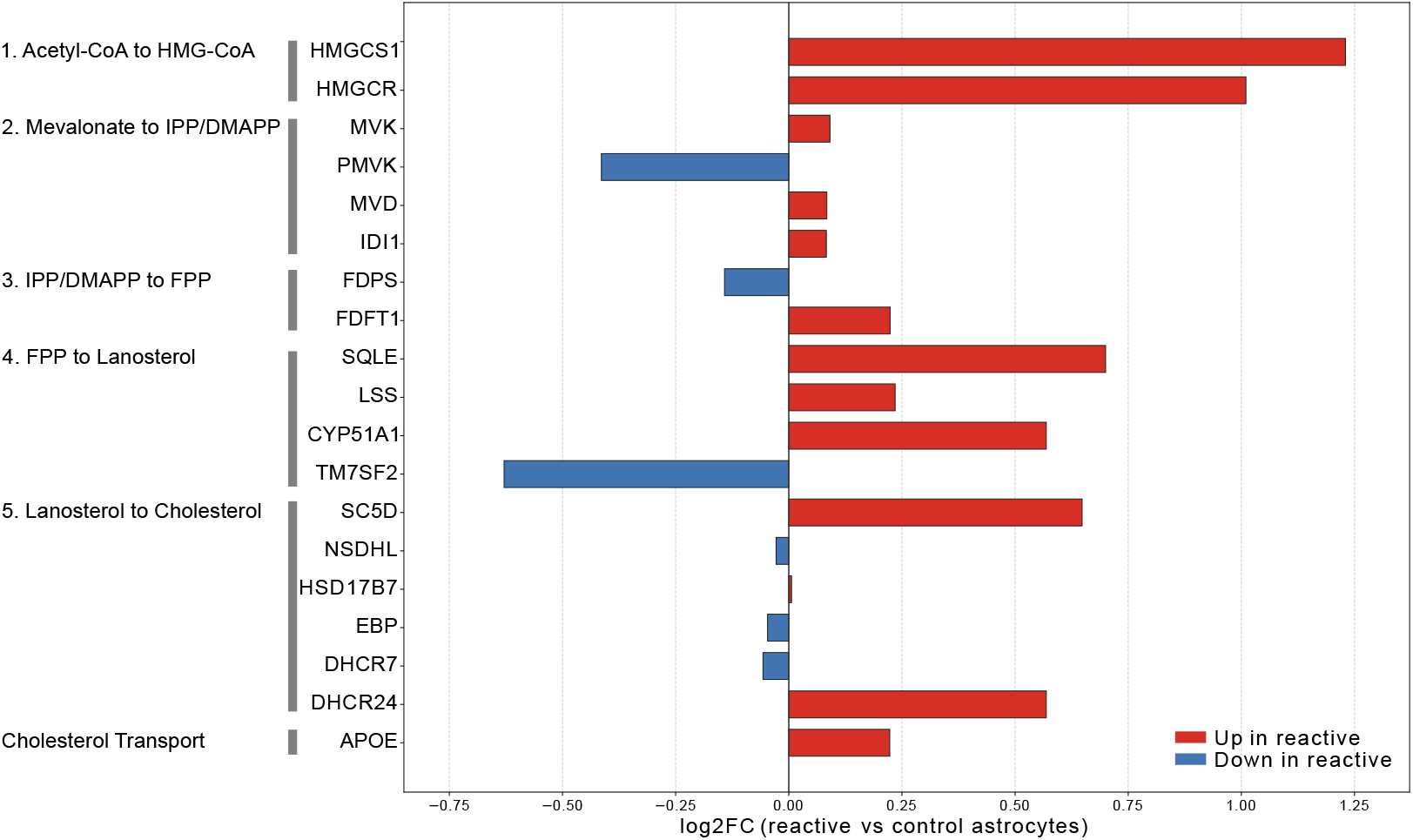
Pro-inflammatory cytokines induce cholesterol biosynthesis proteins in astrocytes. Bar graph showing changes in cholesterol biosynthesis proteins in human iPSC-derived astrocytes following stimulation with TNFα, IL-1α, and C1q. Data were extracted from the proteomics dataset reported by Feringa et al. (2025). Bars represent changes in protein abundance relative to unstimulated controls. Multiple enzymes in the cholesterol biosynthesis pathway were increased following cytokine treatment, including HMGCS1 and HMGCR, which catalyze the initial steps of cholesterol synthesis. Additional enzymes throughout the mevalonate and sterol biosynthetic pathways were similarly elevated. These findings independently support cytokine-induced activation of cholesterol biosynthesis in astrocytes.

**Figure S4.**
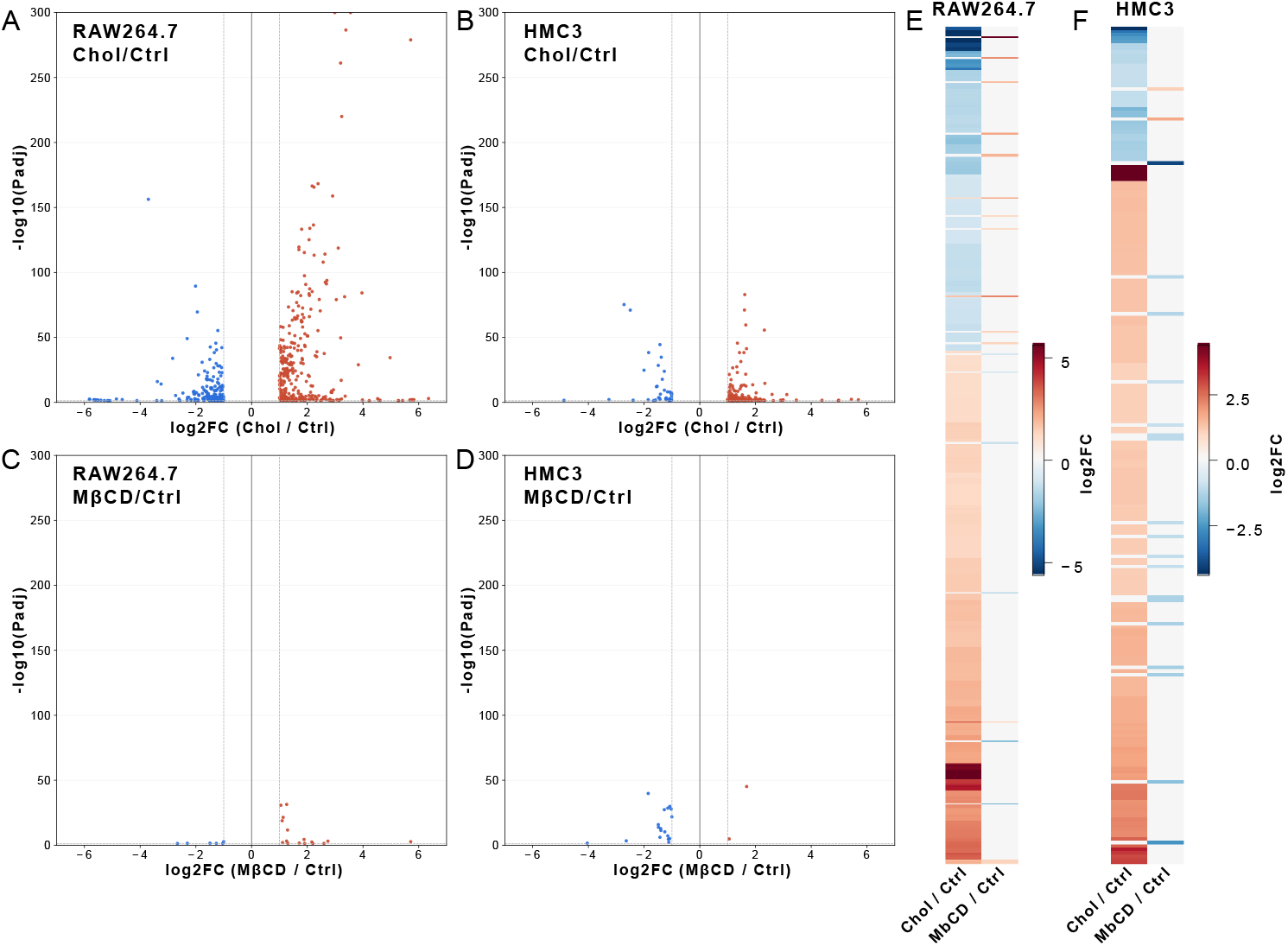
Transcriptomic response to cholesterol and MβCD treatment in RAW264.7 and HMC3 cells. (A,B) RNA-seq volcano plots showing differential gene expression in RAW264.7 macrophages (A) and HMC3 microglia (B) treated with cholesterol relative to untreated controls. (C,D) RNA-seq volcano plots showing differential gene expression in RAW264.7 macrophages (C) and HMC3 microglia (D) treated with MβCD relative to untreated controls, using the same concentration and 2 h treatment employed for cholesterol loading experiments. Significant genes are highlighted in red (upregulated) or blue (downregulated), with nonsignificant genes excluded. Significance was defined as adjusted *P* < 0.05 and |log2 fold change| ≥ 1. (E,F) Hierarchical clustering heatmaps comparing cholesterol- and MβCD-induced transcriptional responses in RAW264.7 macrophages (E) and HMC3 microglia (F). Heatmaps display log2 fold-change values for genes significantly altered in either condition, with red indicating increased expression and blue indicating decreased expression. Distinct clustering patterns indicate that cholesterol loading induces transcriptional programs that are separable from the effects of MβCD treatment alone.

**Figure S5.**
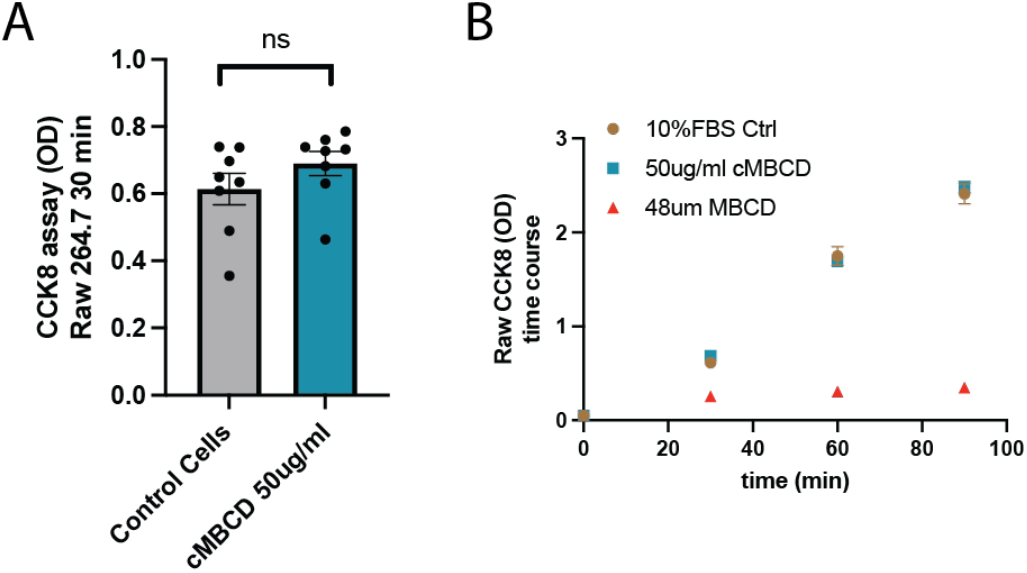
Cholesterol uptake into immune cells does not alter cell viability. RAW 264.7 treated with 50 ug/ml methyl beta cyclodextrin (MβCD) cholesterol (cMβCD) assayed for viability with CCK8. Control cells were treated with fresh 10%FBS with no MβCD cholesterol. The cells were incubated for ~30 min without reagent and then the assay reagent was added and the viability measured at time 30 min or 1hr of cholesterol treatment. (B) A time course of cell viability as in (A). The 90 min time point is a total of 2hrs of cholesterol treatment.

**Figure S6.**
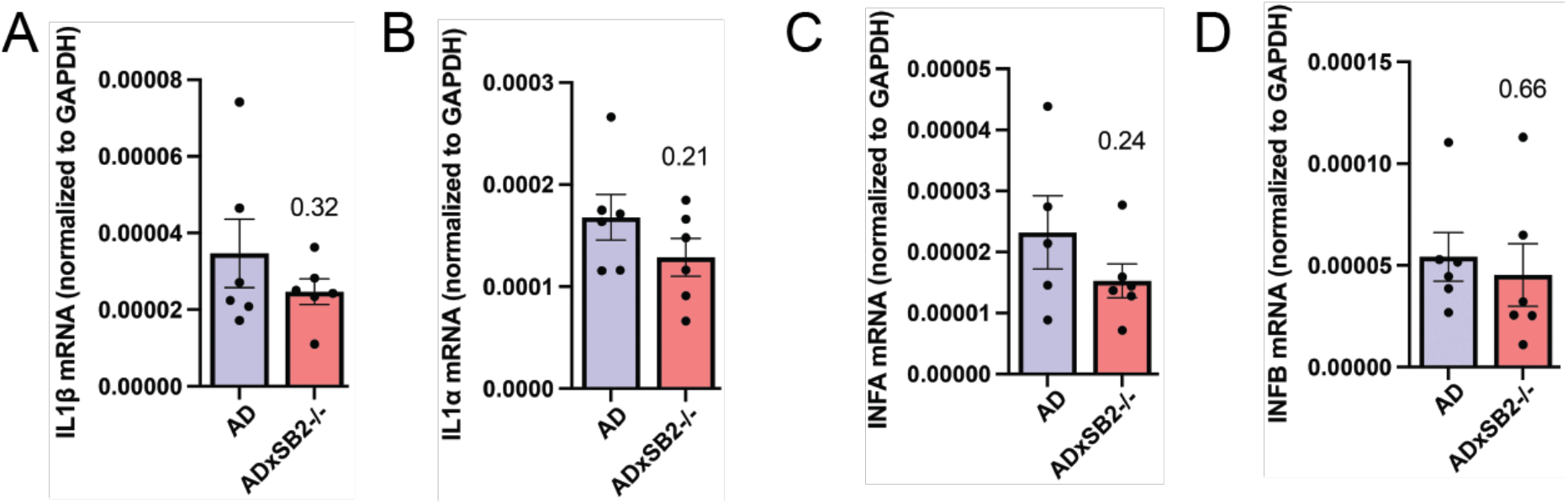
Inflammatory cytokine levels in brain of LPS-treated mice. qPCR of inflammatory cytokines IL1β (A), IL1α (B), INFA (C) and INFB (D) shows a trend of decreased inflammation in SB2−/− like TNFα (see Figure 3E).

**Figure S7.**
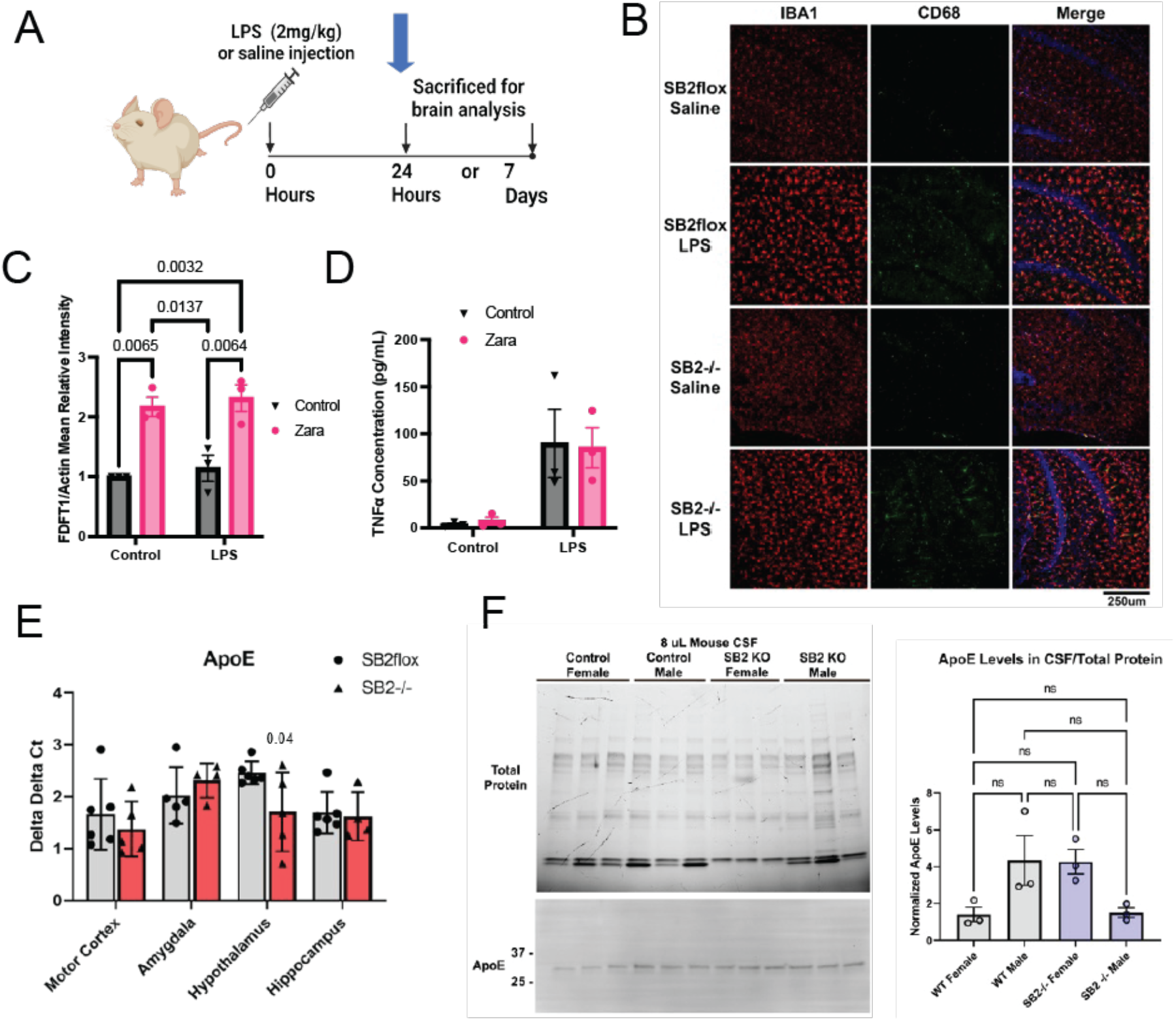
Microglia are responsive to acute stimulation by LPS in the absence of astrocyte cholesterol synthesis. (A) Mice were I.P. injected 1 time with saline or 2mg/kg LPS dissolved in saline. Brains were harvested 24 hours post-injection and examined for CD68 positive IBA1 immunoreactivity to assess microglial activation. (B) Confocal images of CD68 and IBA1 from mouse hippocampus at 24 hrs. (C-D) LPS activation of microglia does not require microglia cholesterol synthesis. SIMA-9 mouse microglia-derived cells were treated with LPS, 250nM zaragozic acid or the combination for 24 hours. Western blot of FDFT1 (squalene synthase) showing that the cholesterol synthesis inhibitor zaragozic acid (zara) efficiently blocks the cholesterol synthesis pathway in the presence or absence of LPS, resulting in compensatory increased FDFT1 protein. (D) Media was collected at the end of 24 hours and TNFα ELISA performed. (E) qPCR was performed on brain homogenate from the regions indicated. Values are reported as the delta Ct with TBP used as a housekeeping gene. There was a small decrease in hypothalamic apoE transcription in the hypothalamus of SB2−/− mice but no difference in other brain regions. N= 4-6 animals per region. (F) CSF was obtained from SBflox (WT) or SB2−/− male and female mice. Each data point is the pooled CSF from 2-3 animals. There was no difference in apoE CSF levels between WT and SB2−/− mice. Comparison made with a Student’s T test in E and ANOVA in F.

**Figure S8.**
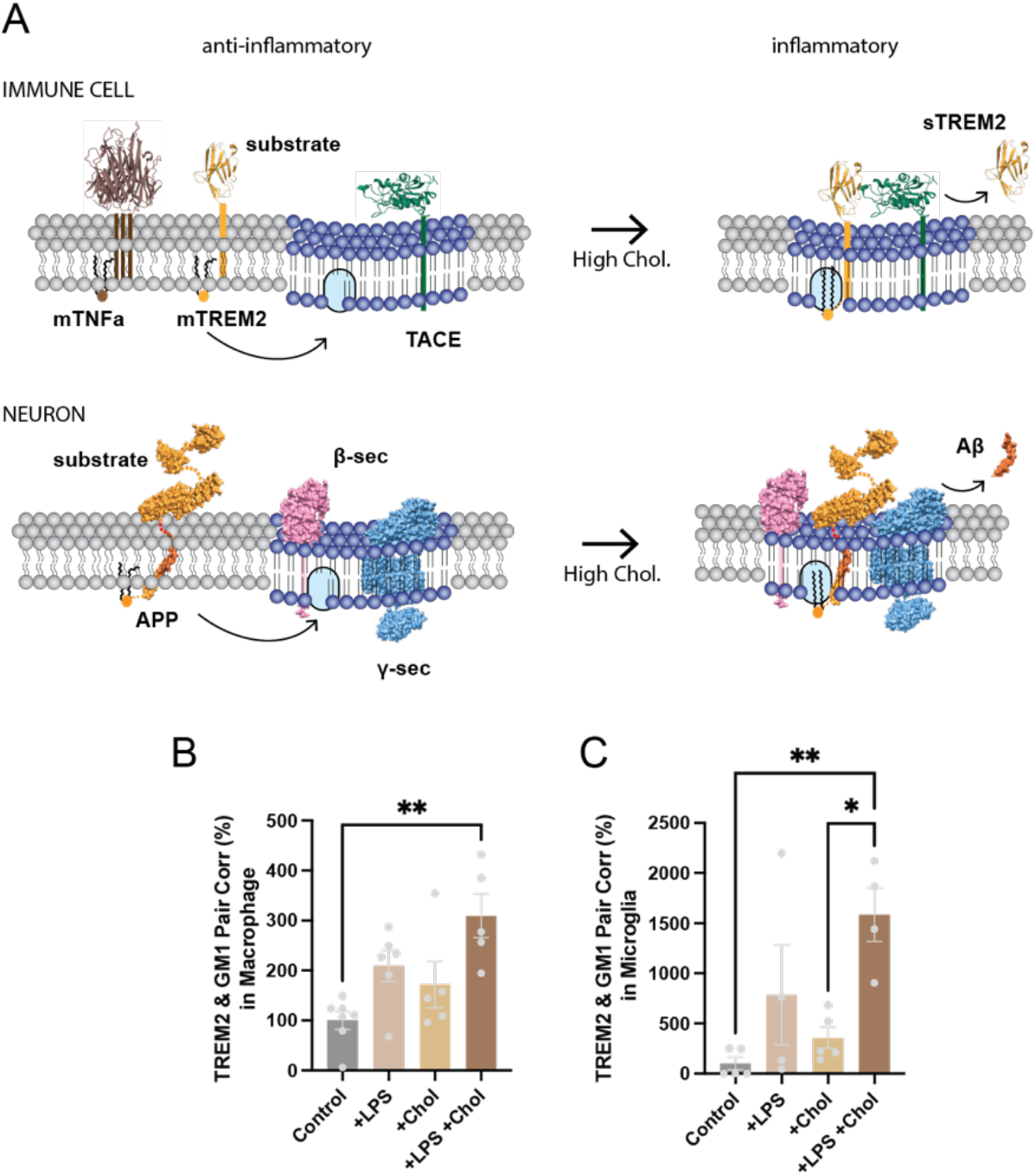
Comparison of molecular mechanisms in neurons and immune cells. (A) Immune cells and neurons have a similar molecular mechanism, but differ in the proteins expressed. The cartoon shows a plasma lipid membrane with regions of disorder (grey lipids with curved acyl chains) and ordered GM1 domains (blue, straight saturated lipids). A palmitate binding site is shown as a light blue shaded oval within the ordered lipids. When cholesterol is high, palmitate (16 carbon saturated lipid often in pairs) binds tightly to the palmitate site. (**top**) TREM2 and TNFα are palmitoylated proteins in immune cells. When cholesterol is low, they resided in disordered region away from ordered GM1 lipids. When cholesterol is high, they shift into GM1 domains where their hydrolytic enzyme ADAM17 (TACE) is located. TREM2 is shown binding to the palmitate binding site where it is cleaved to produce sTREM2, a proinflammatory ectodomain. TNFα is also known to be hydrolyzed by TACE and produce sTNFα (not shown) a proinflammatory cytokine. (**bottom**) The same process occurs in neurons; a palmitoylated amyloid precursor protein (APP) is expressed and resides in disordered lipids. Its hydrolytic enzymes are localized to GM1 lipids. When cholesterol is high the APP binds to the palmitate site where it encounters β- and *γ*-secretases (β-sec and *γ*-sec) and produces the pro-inflammatory amyloid beta (Aβ)^15^. (B-C) dSTORM imaging of cultured immune cells in 8-well chambers. LPS treatment or cholesterol loading alone shows a trend towards increasing TREM2/GM1 lipid cluster association in macrophages and microglia. LPS and cholesterol together has an additive effect. Values are expressed as mean. B-C, each point is a biological replicate (single cell) and comparison are made from samples prepared at the same time. Each experiment was done at least in duplicate. Statistical comparison made with a Student’s T test or ANOVA; *p≤0.05, **p≤0.01, ***p≤0.001, ****p≤0.0001.

## Supplemental Methods

### Animals

Housing, animal care and experimental procedures were consistent with the Guide for the Care and Use of Laboratory Animals and approved by the Institutional Animal Care and Use Committee of the Scripps Research Institute and the University of Virginia. 3xTg-AD mice and B6129SF2/J controls were purchased from Jackson labs. The 3xTg-AD mice were maintained homozygous for all transgenes. SREBP2^flox/flox^ mice (a generous gift of Dr. Jay Horton at UT Southwestern, now available through Jackson labs) were crossed to the hGFAP-Cre as previously described ^24^. These mice were in turn bred to the 3xTg-AD line and crossed back to homozygosity for both the 3xTg-AD transgenes and the SREBP2-flox. 3xTg-AD x SREBP2^flox/flox^ (AD x SB2flox) and 3xTg-AD x SREBP2^flox/flox^ x GFAP-Cre (AD x SB2−/−) are littermates. Only female mice were used for the AD crosses as the male 3xTg-AD mice have a much milder phenotype.

### Cell culture

RAW 264.7 macrophage cells and neuroblastoma 2a (N2a) cells were grown in Dulbecco’s Modified Eagle Medium (DMEM) containing 10% fetal bovine serum (FBS) and 1% penicillin/streptomycin (P/S). Human microglial HMC3 cells were grown in Eagle’s Minimum Essential Medium (EMEM) containing 10% FBS and 1% P/S. SIMA9 cells were grown in Dulbecco’s Modified Eagle Medium: Nutrient Mixture F12 (DMEM/F12) supplemented with 10% FBS, 5% horse serum, and 1% P/S.

Astrocytes were isolated from the cortices of embryonic day 18-21 CD1 mice as described previously^23^. Briefly, cells were dissociated by papain digestion and plated on poly-D-lysine-coated (0.01 mg/mL) tissue culture dish. Dissociated cells were plated in DMEM containing 10% FBS and 1% P/S at 37 °C in 5% CO_2_.

### Astrocyte conditioned media (ACM)

For acute collection of ACM, reactive astrocyte cultures were treated for 1 h with IL-1β (100 ng/mL, Sigma, #SRP8033), TNFα (100 ng/mL, Novus Biologicals, #NBP2-35076), LPS (100 ng/mL, Sigma-Aldrich, #L2018), Human β-Amyloid Peptide (1-42) 100 µg/mL (Biolegend, #932502) or loaded with cholesterol using apoE3 (4 μg/mL, BioVision, #4696-500). Endotoxin assayed to be less than 0.1 ng/µg of apoE3, a concentration non-reactive with microglia. After treatment, astrocytes were incubated with fresh media for 1 h to collect ACM. Microglia and macrophages were incubated with control or reactive ACM for 1 h before cholesterol measurement. Aggregated β-Amyloid peptide was generated by incubating suspended peptide overnight at room temperature. For an elongated collection of ACM, reactive astrocyte cultures were treated for 24 h with either IL-1β (100 ng/ml), IL6 (100 ng/ml, Thermo Fisher Scientific, #PHC0064), and TNFα (100 ng/ml), or LPS (100 ng/mL). After treatment, astrocytes were incubated with fresh media for 24 h to collect ACM. Microglia and macrophages were incubated with control or reactive ACM for 24 h before cholesterol measurement.

### Cholesterol assay

To measure the relative changes in cholesterol level (free cholesterol or total cholesterol), we developed an Amplex Red-based cholesterol assay. Briefly, cells were seeded into 96-well flat culture plates with transparent-bottom to reach confluency (~ 5 x 10^4^ per well). After washing with 200 μL of PBS, cholesterol assay reactions were promptly begun by adding 100 μL of working solution containing 50 μM Amplex red, 1 U/mL horseradish peroxidase, 2 U/mL cholesterol oxidase and 2 U/mL cholesterol esterase in PBS. Relative cholesterol concentration and the background (lacking cells) was determined in triplicate for each sample by measuring fluorescence activity with a fluorescence microplate reader (Tecan Infinite 200 PRO, reading from bottom) for 2 hours at 37°C with excitation wavelength of 530 nm and an emission wavelength of 585 nm. Subsequently, cholesterol level was normalized to the control activity. End point cholesterol signals were then graphed (Mean ± s.e.m.) and statistically analyzed (one-way ANOVA) with GraphPad Prism software (v6.0f).

### Smart-seq protocol

Cholesterol Treatment: Water-soluble cholesterol (MedChemExpress, HY-N0322A-10mg) was prepared as a stock or working solution according to the manufacturer’s instructions. A working concentration of 50 μg/ml soluble cholesterol (with cholesterol comprising 4% of the complex, as a cholesterol-methyl-β-cyclodextrin inclusion complex) was prepared in DMEM (Meilun Bio, MA0212-3). The solution was gently mixed until fully dissolved. Fresh soluble cholesterol-supplemented medium was prepared immediately before use. RAW264.7 murine macrophages and HMC3 human microglia cells were maintained in standard complete culture medium (DMEM supplemented with 10% FBS and 1% penicillin-streptomycin) under standard conditions (37°C, 5% CO_2_). Cells were seeded in appropriate culture plates and grown to 70–80% confluence. Prior to treatment, the regular culture medium was aspirated and replaced as follows: Control group: Fresh regular complete DMEM medium. Cholesterol treatment group (RNA-seq): DMEM containing 50 μg/ml water-soluble cholesterol. Cells were incubated with the respective media for 2 hours at 37°C, 5% CO_2_. Following incubation, both control and treated cells were washed with PBS, detached using trypsin-EDTA. The cell pellets were then resuspended in regular complete culture medium and immediately processed for RNA extraction and subsequent RNA sequencing. All treatments were performed in biological replicates as specified in the main figure legends.

Total RNA was extracted from cells using RNAprep Pure Micro Kit (Tiangen, #DP420). cDNA amplification was performed using the Smart-seq2 protocol with the Discover-sc WTA Kit (#N711, Vazyme). Following reverse transcription and cDNA amplification, 5 ng of amplified cDNA was tagmented using the TruePrep DNA Library Prep Kit V2 for Illumina (Vazyme #TD502) at 55°C for 10 minutes, followed by 12 cycles of indexing PCR. Libraries were then sequenced on Illumina NovaSeq X Plus platform (paired end 150 bp).

### RNA-seq analysis of cholesterol-treated immune cells

RNA-seq analysis was performed on cholesterol-treated versus control RAW264.7 and HMC3 cells. Differential expression results were obtained from the corresponding chol versus control comparisons, and positive log2FC values indicate higher expression in cholesterol-treated cells. For Reactome pathway analysis, differentially expressed genes were filtered using an adjusted p-value (Padj) threshold of < 0.05 and an absolute fold-change threshold of |log2FC| >= 1. Reactome pathway enrichment was then performed separately for each cell line using the Reactome_2022 gene-set collection, and the top 20 enriched pathways were visualized.

To examine inflammatory signaling in greater detail, genes representing major inflammatory pathways were manually curated and grouped into the following pathway sets. The JAK/STAT set included Jak1, Jak3, Stat1, Stat2, and Stat3. The MAPK/AP-1 set included Mapk1, Mapk3, Jun, Junb, Jund, Fos, Fosb, Fosl1, Fosl2, Atf2, and Atf3. The NF-kB set included Nfkb1, Nfkb2, Rela, Relb, and Rel. The Interferon Regulatory Factors set included Irf1, Irf3, Irf5, Irf7, and Irf8. The Interferon-Stimulated Genes (ISGs) set included Isg15, Ifit1, Ifit2, Ifit3, Ifih1, Mx1, Mx2, Oas1a, Oas1g, Oas2, Oas3, and Rsad2. The Suppressor of Cytokine Signaling set included Socs1, Socs2, and Socs3. Gene-level log2FC values for these curated pathway genes were extracted from the cholesterol-versus-control comparisons and displayed in pathway-organized heatmaps. Principal component analysis (PCA) was then performed separately for RAW264.7 and HMC3 using the transcriptional profiles of these selected inflammatory pathway genes.

To investigate the strongest shared Reactome hit, genes contributing to the enriched pathway Response of EIF2AK1 (HRI) To Heme Deficiency were extracted directly from the Reactome output and visualized using bubble plots, where the x-axis represents log2FC and bubble size represents adjusted p-value. Because these genes suggested a strong stress-response signature, stress-associated genes were further curated into six functional subpathways. The PERK/ISR/ATF4-CHOP set included Eif2ak3, Atf4, Ddit3, Trib3, Chac1, Ppp1r15a, Atf3, and Herpud1. The Heat Shock/Proteostasis set included Hspa1a, Hspa1b, and Serpinh1. The IRE1/XBP1 set included Ern1, Xbp1, and Herpud1. The Oxidative/TXNIP set included Txnip and Hmox1. The Stress MAPK/AP-1 set included Map3k8, Mapk1, Mapk3, Jun, Junb, Jund, Fos, Fosb, Fosl1, Fosl2, Atf2, Atf3, Dusp8, and Dusp10. The Inflammasome Link set included Txnip, Pycard, Nlrp3, Casp1, Casp4, Il1a, Il1b, and Nos2. For each subpathway, the mean gene log2FC and the number of genes upregulated by cholesterol were summarized separately for RAW264.7 and HMC3.

To visualize genes linking stress signaling and inflammatory activation, a focused bridge-gene set was also curated. Stress-core genes included Ddit3, Trib3, Chac1, Ppp1r15a, Atf4, and Herpud1. Stress-to-inflammation bridge genes included Atf3, Jun, Dusp8, Dusp10, Map3k8, and Txnip. Volcano plots were generated from the full differential expression tables to highlight these stress-related genes in the context of the whole transcriptome. All RNA-seq visualizations and downstream analyses were performed in Python using standard scientific computing and plotting libraries.

### LPS Injections

LPS was injected intraperitoneally in ~15-week-old male mice with 1 dose of 2 mg/kg LPS in saline or vehicle control. LPS was purchased from Sigma Aldrich (L3024) and the same lot of LPS was used for all animals tested. LPS was diluted into sterile saline solution at a concentration of 5 mg/mL. Mice were monitored throughout the course of the experiment and provided with extra cotton nest bedding. Sickness behavior in animals was evident at 24 hours and improved over the course of 5 days.

### dSTORM Super-resolution imaging

#### Fixed cell preparation

Cells were grown to 60% confluence. Cells were rinsed with PBS and then fixed with 3% paraformaldehyde and 0.1% glutaraldehyde for 15 min to fix both proteins and lipids. Fixative chemicals were reduced by incubating with 0.1% NaBH_4_ for 7 min with shaking followed by three times 10 min washes with PBS. Cells were permeabilized with 0.2% Triton X-100 for 15 min and then blocked with a standard blocking buffer (10% bovine serum albumin (BSA) / 0.05% Triton in PBS) for 90 min at room temperature. For labelling, cells were incubated with primary antibody (TLR4: Thermo Fisher Scientific, #482300; TREM2: R&D Systems, #MAB17291; IFNγR1: Thermo Fisher Scientific, #10808-1-AP) for 60 min in 5% BSA / 0.05% Triton / PBS at room temperature followed by 5 washes with 1% BSA / 0.05% Triton / PBS for 15 min each. Secondary antibody was added in the same buffer as primary for 30 min at room temperature followed by 5 washes as stated above. Cells were then washed with PBS for 5 min. Cell labelling and washing steps were performed while shaking. Labeled cells were then post-fixed with fixing solution, as above, for 10 min without shaking followed by three 5 min washes with PBS and two 3 min washes with deionized distilled water. The specificity of the antibodies used in dSTORM imaging were validated by western blots (see manufacture product page). Specifically, TREM2 antibody stains TREM2 transfectants but not TREM1 transfectants in flow cytometry.

#### Brain slice preparation

Mouse brain slicing and staining were performed as previously described^45^ with minor modifications. Mouse brains were fixed in 4% paraformaldehyde, incubated in a 20% sucrose/PBS solution at 4 °C for 3 days, and embedded in Tissue-Tek OCT compound (Sakura). Sagittal sections (50 μm) were collected and placed into 24-well plate wells containing PBS. Fixative chemicals were reduced by incubating with 0.1% NaBH_4_ for 30 min while gently shaking at room temperature followed by three times 10 min washes with PBS. Samples were permeabilized with 0.2% Triton X-100 for 2 hours and then blocked with a standard blocking buffer (10% bovine serum albumin (BSA) / 0.05% Triton in PBS) for 6 hours at room temperature. For labelling, samples were incubated with primary antibody for 3 hours in 5% BSA / 0.05% Triton / PBS at room temperature then 3 days at 4 °C followed by 5 washes with 1% BSA / 0.05% Triton / PBS for 1 hour each. Secondary antibody was added in the same buffer as primary for 3 days at 4 °C followed by 5 washes as stated above. Sample labelling and washing steps were performed while shaking. Labeled brain tissues were then post-fixed with fixing solution, as above, for 1 hour without shaking followed by three 30 min washes with PBS and two 30 min washes with deionized distilled water. Brain slices were mounted onto the stainless-steel imaging chamber (Life Sciences, #A-7816) with #1 thickness cover glass (Chemglass, # CLS-1760-025), and 2% agarose were pipetted onto the slice to form a permeable agarose pad and prevent sample movement during imaging.

#### Imaging protocol

Images were recorded with a Bruker Vutara 352 or VXL with a 60X Olympus Silicone objective. Frames with an exposure time of 20 ms were collected for each acquisition. Excitation of the Alexa Fluor 647 dye was achieved using 640 nm lasers and Cy3B was achieved using 561 nm lasers. Laser power was set to provide isolated blinking of individual fluorophores. Cells were imaged in a photo-switching buffer comprising of 1% β-mercaptoethanol (Sigma, #63689), oxygen scavengers (glucose oxidase (Sigma, #G2133) and catalase (Sigma, #C40)) in 50mM Tris (Affymetrix, #22638100) + 10mM NaCl (Sigma, #S7653) + 10% glucose (Sigma, #G8270) at pH 8.0. Axial sample drift was corrected during acquisition through the Vutara 352’s vFocus system.

Images were constructed using the default modules in the Zen software. Each detected event was fitted to a 2D Gaussian distribution to determine the center of each point spread function plus the localization precision. The Zen software also has many rendering options including removing localization errors and outliers based on brightness and size of fluorescent signals. Pair correlation and cluster analysis was performed using the Statistical Analysis package in the Vutara SRX software. Pair Correlation analysis is a statistical method used to determine the strength of correlation between two objects by counting the number of points of probe 2 within a certain donut-radius of each point of probe 1. This allows for localization to be determined without overlapping pixels as done in traditional diffraction-limited microscopy. Cluster size estimation and cluster density were calculated through cluster analysis by measuring the length and density of the clusters comprising of more than 10 particles with a maximum particle distance of 0.1 µm.

### Quantitation of Astrocyte Morphology and Microglia CD68

Astrocyte cell 3D reconstructions were quantified using IMARIS filament package software. Mouse brain hemispheres were collected from PBS perfused 40-week-old animals. Tissue was cut into 40 µm sections and immunostained using anti-GFAP antibody (Mab360, EMD Millipore) applied overnight at a 1:500 concentration in an immunostaining blocking solution of 5% BSA, 5% horse serum, 0.1% Triton X-100 in PBS. Alexa 647 secondary antibody was applied for fluorescent visualization. Astrocyte FDFT1 was immunostained using Abcam antibody Ab195046 at a 1:200 dilution. Confocal microscopy Z-stack images of astrocyte populations were collected using a Leica TCS SP8 confocal microscope from the CA1 region of the hippocampus (30 µm depth, 0.5 µm steps, x 20 magnification) of each animal. Raw .tif files were then used for IMARIS software analysis (Oxford Instruments). IMARIS was used to reconstruct the astrocyte (GFAP) surfaces. A threshold of 20/250 was applied to each image before analysis. The reconstruction was used for filament reconstruction using a max diameter of 10.0 µm with seed points at 0.200 µm and a diameter of sphere regions set to 15 µm. Surface and filament parameters from each astrocyte cell population were then exported to excel for statistical analysis in PRISM software. All images taken for analysis were collected in a single imaging session using the same parameters including pinhole, laser intensity and gain.

For determining the CD68 content of hippocampal microglia, 40 µm PFA fixed brain sections were immunostained by applying an IBA1 antibody (E4O4W, Cell Signaling) at a 1:500 dilution and a CD68 antibody (FA-11, Bio Rad) at a 1:500 dilution in blocking solution (described above) overnight at 4 degrees. Alexa Fluor 647 anti-rabbit and Alexa Fluor 555 anti-rat secondary antibodies were used for visualization. Confocal Z-stack images of hippocampal IBA1+ cell populations were collected from each brain section (~36 µm depth, 0.5 µm steps, x 20 magnification) using a Leica Stellaris 8 confocal microscope. Raw .tif files were imported into IMARIS software to perform 3D reconstructions. IBA1 and CD68 were set with a threshold cutoff of 20/250. The colocalization function was then used to determine the % colocalization in the region of interest as well as the % of IBA1 area occupied with CD68. 3 sections from each individual animal were imaged, analyzed and averaged to generate each individual datapoint for statistical analysis.

### Filipin Staining

Filipin (Sigma F-9765) 25 mg/ml in DMSO was diluted to 0.05 mg/ml in PBS/10% FBS. Brain slices fixed in 4% PFA were rinsed 3x with PBS and incubated with 1 ml of 1.5 mg glycine/ml PBS for 10 min at room temperature. The slices were stained with 1 mL of 0.05 mg/ml filipin working solution for 2h at room temp with slow shaking. The slices were rinsed 3x with PBS and imaged on a FV-1000 Olympus confocal with 20x objective.

### ELISA

The ELISA plate (Invitrogen 44-2404-21) was coated with 100 µL anti-mouse TNFα antibody (capture antibody, BioLegend #510802) diluted 1:250 in coating buffer (BioLegend #421701) and incubated overnight at 4°C. The plate was washed with 200 µL PBS for three times, blocked with 100 µL blocking buffer (PBS with 10%BSA and 0.05% Triton X-100), and incubated for 1 h. Then the blocking buffer was removed from each well and 50 µL supernatant was added and incubated for 1 h. Next, 50 µL Biotin anti-mouse TNFα antibody (primary antibody, BioLegend #506312) diluted 1:1000 in PBST buffer (PBS with 0.01% Triton X-100) was mixed with the supernatant and incubated for 3 h. The plate was washed with 200 µL PBST for four times and incubated with 80 µL HRP streptavidin (BioLegend #405210) diluted 1:1000 in PBST for 1 h in the dark. After that, the plate was washed with 200 µL PBST for four times and incubated with 80 µL Chromogen (Invitrogen #002023) for 30 min in the dark. The substrate development was terminated by 80 µL stop solution (Invitrogen #SS04). Relative TNFα concentration was determined by measurement of absorbance at 450nm on a microplate reader (Tecan Infinite 200 PRO).

For TNFα levels in Fig S6D were determined using a mouse TNF *α* ELISA kit (R&D Systems, DY410-05, Minneapolis, MN, USA) according to manufacturer’s instructions. All samples were run in technical duplicates. Absorbance was measured at 450 nm with a background subtraction reading at 570 nm for normalization.

### qPCR on Brain Tissue

RNA was extracted from 60-week-old female 3xTg and 3xTgxSB2−/− frontal cortex using the Isolate II RNA Mini Kit (BIO-52073) from Bioline per manufacturer’s instructions. cDNA synthesis was performed using the iScript cDNA synthesis kit (#1708841) from Bio Rad using a SimpliAmp thermocycler following iScript manufacturer’s instructions with 1 μg input RNA. qPCR was performed by standard protocols using the following TaqMan primers: GAPDH Mm99999915_g1, TNFα Mm00443258_m1, NFKb Mm00476361_m1, IL1b Mm00434228_m1, IL1a Mm00439620_m1, INFa Mm00833961_s1, and INFb Mm00439552_s1. GAPDH served as a loading control for normalization and the relative abundance of transcripts was determined using the 2^- (Target Cq-GAPDH Cq) formula.

### Proteomics analysis

#### Preparation of samples for LC-MS analysis

Samples were prepared following the Solid-Phase-Extraction Capture (SPEC) protocol described previously^46^. All liquid-handling steps were performed using an Agilent Bravo pipetting robot. Cells were lysed directly in the cell culture plate by adding 10 µl oflysis buffer containing 2% w/v sodium deoxycholate, 100 mM Tris-HCl pH 8.5, 10 mM tris(2-carboxyethyl)phosphine, 40 mM chloroacetamide, and 0.1 n-Dodecyl-β-D-maltopyranoside (DDM), and frozen at −80 °C until further processing. Before sample preparation, samples were heated to 60 °C for 10 min. In-house synthesized DNase from Serratia marcescens was then added to a final concentration of 1 U/µl together with MgCl2 to a final Mg2+ concentration of 1 mM and incubated for 15 min at room temperature and constant shaking at 300 rpm on a ThermoBlock. SPEC tips we prepared in-house by placing one plug of strong-anion-exchange (SAX) material (3M Empore) in a pipette tip with a blunt-end syringe needle. Tips were activated using 10 µl DMSO and centrifuged at 700g for 1.5 min. Next, tips were preequilibrated with 20 µl of SPEC preequlibration buffer (20 mM 3-(Cyclohexylamino)-1-propanesulfonic acid (CAPS) pH 10.5, 0.01% DDM in 50% methanol) by centrifugation at 700g for 2 min. Then, 200 ng of protein was loaded onto SPEC tips in SPEC equilibration at a 1:10 (v/v) ratio and centrifuged at 200g for 10 min. The tips were then washed using 20 µl SPEC wash buffer (50 mM triethylammonium bicarbonate (TEAB), 0.01% DDM) by centrifuging at 700g for 3 min or until all liquid has passed through the tip. Digestion was performed by applying 5 µl of digestion buffer (25 ng/ µl trypsin, 25 ng/ µl LysC, 50 mM TEAB, 0.01% DDM) into the tip, centrifuging at 100g for 20s and incubating at 37 °C for 1 h. For peptide loading, C18 Evotips Pure tips were washed with 50 µl buffer B (99.9% acetonitrile, 0.1% formic acid), centrifuged at 700g for 1 min, activated with 2-propanol for 1 min, and equilibrated with 50 µl buffer A (0.1% formic acid in water) prior to loading. Digested peptides were then eluted from SPEC tips and simultaneously loaded onto Evotips in 100 µl of buffer A by centrifuging at 300g for 5 min. Evotips were washed with 75 µl of buffer A by centrifugation at 700g for 1 min. 150 µl of buffer A was finally added to Evotips before storage at 4 °C until further use.

#### LC-MS/MS analysis

All samples were analyzed using an Evosep Eno LC system (Evosep) coupled to an Orbitrap Astral Zoom mass spectrometer (Thermo Fisher Scientific). Peptides were eluted from the Evotips using a LC gradient with a throughput of 500 samples per day on PepSep column of 4-cm length, 150-µm-internal diameter, packed with 1.9 µm C18 beads (Evosep). Column was integrated with 30-µm stainless steel emitter (Evosep). The column temperature was maintained at 60 °C using a column heater (Evosep). The Orbitrap Astral Zoom was equipped with an EASY-Spray source (Thermo Fisher Scientific). For ABCA, IFNG, IFNB1 and their respective controls, Orbitrap Astral Zoom was equipped with a FAIMS Pro interface (−40 V compensation voltage, 3.5 L/min carrier gas, Thermo Fisher Scientific). FAIMS inner electrode temperature was set to 100 °C and outer electrode temperature was set to 80 °C. For Simavastin and the Simavastin Controls, FAIMS was not installed. For all samples, sample acquisition was performed in DIA mode. An electrospray voltage of 1,900 V was applied for ionization, and the radio frequency level was set to 40. Orbitrap MS1 spectra were aquired from 380 to 980 m/z at a resolution of 240,000 (at m/z 200) with a normalized automated gain control (AGC) target at 500% and a maximum injection time of 3 ms. For the Astral MS/MS scans in data-independent acquisition (DIA) mode, 100 variable isolation windows, designed with a pyDIAid software^47^, were used (Supplementary Table 1). A maximum injection time of 5 ms was used. The isolated ions were fragmented using high-energy collisional dissociation with 25% normalized collision energy. Samples were acquired in randomized order.

#### LC-MS/MS data search

All raw files were processed together in a single DIA-NN (v2.5.1) analysis^48^. Spectra were searched against a predicted spectral library generated in silico from a FASTA database of reviewed canonical sequences and isoforms of Homo sapiens (taxon ID 9606; 42,516 entries; UniProtKB, 26 July 2025); the same FASTA was additionally supplied for library reannotation and protein inference. The predicted library was generated allowing up to one missed cleavage, with cysteine carbamidomethylation as a fixed modification and methionine oxidation as a variable modification, and with peptide length restricted to 7–55 amino acids, precursor charge state to 2–7, precursor m/z to 300– 1,000, and fragment ion m/z to 200–1,800. MS1 and MS2 mass tolerances were fixed at 6 and 10 ppm, respectively, and the scan-window radius was set to 6. Match-between-runs and peptidoform scoring were enabled. Precursor- and protein-group-level output was filtered at 1% FDR.

#### Bioinformatics analysis

Protein quantification was taken from the DIA-NN parquet output. For downstream analysis, we retained only decoy-free, proteotypic entries and applied the following DIA-NN quality filters: Q.Value <= 0.01, Lib.Q.Value <= 0.01, Lib.PG.Q.Value <= 0.01, Protein.Q.Value <= 0.01, and PG.Q.Value <= 0.01 when available. Entries with missing or non-positive Genes.MaxLFQ values were removed. Gene-level abundance matrices were assembled from Genes.MaxLFQ values and log2-transformed. Differential expression was calculated in Python using Welch’s two-sided t-test (scipy.stats.ttest_ind, equal_var=False) for each gene, comparing treatment and control triplicates for each contrast. The four comparisons analyzed were simvastatin versus control, ABCA1 CRISPRko versus NT3/4, IFNG CRISPRa versus NTCRISPRa, and IFNB1 CRISPRa versus NTCRISPRa. P values were adjusted across all tested genes using the Benjamini-Hochberg false discovery rate procedure. We considered genes significant at adjusted P < 0.05. No additional absolute log2 fold-change cutoff was applied. For volcano plots, log2 fold change was plotted on the x-axis and −log10(raw P value) on the y-axis. Significant hits were highlighted using the Benjamini-Hochberg adjusted q values. We did not impose a minimum peptide-count filter, which avoids removing on/off hits that can be recovered through match-between-runs. The complete proteomics results are provided in Supplementary Table 2.

The mass spectrometry proteomics data have been deposited to the ProteomeXchange Consortium via the PRIDE partner repository with the dataset identifier PXD080432. The reviewer access token is dVl96MxiVIGg^49^.

### Antibodies

Antibodies were diluted in blocking solution containing 5% BSA. FDFT1 antibody (Abcam, ab195046, Cambridge, UK) was used at a 1:1000 dilution and secondary donkey anti-rabbit AlexaFluor 647 Plus (Invitrogen, A32795, Waltham, MA, USA) was used at a 1:5000 dilution. Rhodamine-conjugated actin antibody (BioRad, 12004163, Hercules, CA, USA) was used at a dilution of 1:5000.

### Statistical analysis

Statistical analysis was performed using GraphPad Prism 10 software (GraphPad Software, La Jolla, CA, USA). Statistical significance between the treatment groups was determined by Student’s T test or Tukey’s post-hoc test for multiple comparison after a one-way ANOVA.

## References

1. Shi, F. D. & Yong, V. W. Neuroinflammation across neurological diseases. Science vol. 388 Preprint at 10.1126/science.adx0043 (2025).

2. Heneka, M. T. et al. Neuroinflammation in Alzheimer disease. Nature Reviews Immunology vol. 25 321–352 Preprint at 10.1038/s41577-024-01104-7 (2025).

3. DiSabato, D. J., Quan, N. & Godbout, J. P. Neuroinflammation: the devil is in the details. J. Neurochem. 139, 136–153 (2016).

4. Tall, A. R. & Yvan-Charvet, L. Cholesterol, inflammation and innate immunity. Nat. Rev. Immunol. 15, 104–116 (2015).

5. Leng, F. & Edison, P. Neuroinflammation and microglial activation in Alzheimer disease: where do we go from here? Nat. Rev. Neurol. 17, 157–172 (2021).

6. Calsolaro, V. & Edison, P. Neuroinflammation in Alzheimer’s disease: Current evidence and future directions. Alzheimer’s and Dementia 12, 719–732 (2016).

7. Matejuk, A. & Ransohoff, R. M. Crosstalk Between Astrocytes and Microglia: An Overview. Front. Immunol. 11, 1–11 (2020).

8. Jha, M. K., Jo, M., Kim, J. H. & Suk, K. Microglia-Astrocyte Crosstalk: An Intimate Molecular Conversation. Neuroscientist 25, 227–240 (2019).

9. Liddelow, S. A. et al. Neurotoxic reactive astrocytes are induced by activated microglia. Nature 541, 481–487 (2017).

10. Wang, C. et al. Selective removal of astrocytic APOE4 strongly protects against tau-mediated neurodegeneration and decreases synaptic phagocytosis by microglia. Neuron 1–18 (2021) doi:10.1016/j.neuron.2021.03.024.

11. Martín, M. G., Pfrieger, F. & Dotti, C. G. Cholesterol in brain disease: sometimes determinant and frequently implicated. EMBO Rep. 15, 1036–1052 (2014).

12. Hansen, S. B. & Wang, H. The shared role of cholesterol in neuronal and peripheral inflammation. Pharmacol. Ther. 249, 108486 (2023).

13. Guttenplan, K. A. et al. Neurotoxic reactive astrocytes induce cell death via saturated lipids. Nature https://doi.org/10.1038/s41586-021-03960-y (2021) doi:10.1038/s41586-021-03960-y.

14. Kim, S. & Son, Y. Astrocytes Stimulate Microglial Proliferation and M2 Polarization In Vitro through Crosstalk between Astrocytes and Microglia. International Journal of Molecular Sciences Article https://doi.org/10.3390/ijms (2021) doi:10.3390/ijms.

15. Wang, H. et al. Regulation of beta-amyloid production in neurons by astrocyte-derived cholesterol. Proc. Natl. Acad. Sci. U. S. A. 118, e2102191118 (2021).

16. Feingold, K. R. et al. Multiple cytokines stimulate hepatic lipid synthesis in vivo. Endocrinology 125, 267–74 (1989).

17. Petersen, E. N., Chung, H.-W., Nayebosadri, A. & Hansen, S. B. Kinetic disruption of lipid rafts is a mechanosensor for phospholipase D. Nat. Commun. 7, 13873 (2016).

18. Feringa, F. M. et al. The Neurolipid Atlas: a lipidomics resource for neurodegenerative diseases. Nat. Metab. 7, 2142–2164 (2025).

19. Ma, T. et al. Suppression of eIF2α kinases alleviates Alzheimer’s disease-related plasticity and memory deficits. Nat. Neurosci. 16, 1299–1305 (2013).

20. Muñoz Herrera, O. M. & Zivkovic, A. M. Microglia and Cholesterol Handling: Implications for Alzheimer’s Disease. Biomedicines vol. 10 Preprint at 10.3390/biomedicines10123105 (2022).

21. Kong, F. J., Wu, J. H., Sun, S. Y. & Zhou, J. Q. The endoplasmic reticulum stress/autophagy pathway is involved in cholesterol-induced pancreatic β-cell injury. Sci. Rep. 7, (2017).

22. Oddo, S. et al. Triple-transgenic model of Alzheimer’s disease with plaques and tangles: intracellular Abeta and synaptic dysfunction. Neuron 39, 409–421 (2003).

23. Wang, H. et al. Regulation of beta-amyloid production in neurons by astrocyte-derived cholesterol. Proceedings of the National Academy of Sciences 118, e2102191118 (2021).

24. Ferris, H. A. et al. Loss of astrocyte cholesterol synthesis disrupts neuronal function and alters whole-body metabolism. Proc. Natl. Acad. Sci. U. S. A. 114, 1189–1194 (2017).

25. Nagamoto-Combs, K., McNeal, D. W., Morecraft, R. J. & Combs, C. K. Prolonged microgliosis in the rhesus monkey central nervous system after traumatic brain injury. J. Neurotrauma 24, 1719–1742 (2007).

26. Owlett, L. D. et al. Gas6 induces inflammation and reduces plaque burden but worsens behavior in a sex-dependent manner in the APP/PS1 model of Alzheimer’s disease. J. Neuroinflammation 19, 1–17 (2022).

27. Hoogland, I. C. M., Houbolt, C., van Westerloo, D. J., van Gool, W. A. & van de Beek, D. Systemic inflammation and microglial activation: Systematic review of animal experiments. J. Neuroinflammation 12, 1–13 (2015).

28. Zhao, J. et al. Neuroinflammation induced by lipopolysaccharide causes cognitive impairment in mice. Sci. Rep. 9, (2019).

29. Rudge, J. D. A. The Lipid Invasion Model: Growing Evidence for This New Explanation of Alzheimer’s Disease. Journal of Alzheimer’s Disease 94, 457–470 (2023).

30. Sawada, M., Kondo, N., Suzumura, A. & Marunouchi, T. Production of tumor necrosis factor-alpha by microglia and astrocytes in culture. Brain Res. 491, 394–7 (1989).

31. Duan, Y. et al. Regulation of cholesterol homeostasis in health and diseases: from mechanisms to targeted therapeutics. Signal Transduction and Targeted Therapy vol. 7 Preprint at 10.1038/s41392-022-01125-5 (2022).

32. Miller, Y. I., Navia-Pelaez, J. M., Corr, M. & Yaksh, T. L. Lipid rafts in glial cells: role in neuroinflammation and pain processing. J. Lipid Res. 61, 655–666 (2020).

33. Poggi, M. et al. Palmitoylation of TNF alpha is involved in the regulation of TNF receptor 1 signalling. Biochim. Biophys. Acta Mol. Cell Res. 1833, 602–612 (2013).

34. Robinson, C. V., Rohacs, T. & Hansen, S. B. Tools for Understanding Nanoscale Lipid Regulation of Ion Channels. Trends Biochem. Sci. 44, 795–806 (2019).

35. Pavel, M. A., Petersen, E. N., Wang, H., Lerner, R. A. & Hansen, S. B. Studies on the mechanism of general anesthesia. Proc. Natl. Acad. Sci. U. S. A. 117, 13757–13766 (2020).

36. Petersen, E. N. et al. Mechanical activation of TWIK-related potassium channel by nanoscopic movement and rapid second messenger signaling. Elife 12, (2024).

37. Stelzmann, R. A., Norman Schnitzlein, H. & Reed Murtagh, F. An english translation of alzheimer’s 1907 paper, “über eine eigenartige erkankung der hirnrinde”. Clinical Anatomy 8, 429–431 (1995).

38. Farmer, B. C., Walsh, A. E., Kluemper, J. C. & Johnson, L. A. Lipid Droplets in Neurodegenerative Disorders. Front. Neurosci. 14, 1–13 (2020).

39. Zacharias, D. A., Violin, J. D., Newton, A. C. & Tsien, R. Y. Partitioning of lipid-modified monomeric GFPs into membrane microdomains of live cells. Science 296, 913–6 (2002).

40. Petersen, E. N., Pavel, M. A., Wang, H. & Hansen, S. B. Disruption of palmitate-mediated localization; a shared pathway of force and anesthetic activation of TREK-1 channels. Biochim. Biophys. Acta Biomembr. 1862, 183091 (2020).

41. Lang, T. et al. SNAREs are concentrated in cholesterol-dependent clusters that define docking and fusion sites for exocytosis. EMBO Journal vol. 20 2202–2213 (2001).

42. Petersen, E. N., Chung, H.-W., Nayebosadri, A. & Hansen, S. B. Kinetic disruption of lipid rafts is a mechanosensor for phospholipase D. Nat. Commun. 7, 13873 (2016).

43. Tellier, E. et al. The shedding activity of ADAM17 is sequestered in lipid rafts. Exp. Cell Res. 312, 3969–3980 (2006).

44. Levental, I., Lingwood, D., Grzybek, M., Coskun, U. & Simons, K. Palmitoylation regulates raft affinity for the majority of integral raft proteins. Proceedings of the National Academy of Sciences 107, 22050–22054 (2010).

45. German, C. L., Gudheti, M. V., Fleckenstein, A. E. & Jorgensen, E. M. Brain slice staining and preparation for three-dimensional super-resolution microscopy. in Methods in Molecular Biology (2017). doi:10.1007/978-1-4939-7265-4_13.

46. Heymann, T. et al. Solid-Phase Extraction Capture (SPEC) in nanoliter volumes for fast, robust and ultra-sensitive proteomics. Preprint at 10.1101/2025.05.31.657165 (2025).

47. Skowronek, P., Wallmann, G., Wahle, M., Willems, S. & Mann, M. An accessible workflow for high-sensitivity proteomics using parallel accumulation–serial fragmentation (PASEF). Nat. Protoc. 20, 1700–1729 (2025).

48. Demichev, V., Messner, C. B., Vernardis, S. I., Lilley, K. S. & Ralser, M. DIA-NN: neural networks and interference correction enable deep proteome coverage in high throughput. Nat. Methods 17, 41–44 (2020).

49. Perez-Riverol, Y. et al. The PRIDE database at 20 years: 2025 update. Nucleic Acids Res. 53, D543–D553 (2025).

